# Functional interdependence of the actin nucleator Cobl and Cobl-like in dendritic arbor development

**DOI:** 10.1101/2021.03.03.433715

**Authors:** Maryam Izadi, Eric Seemann, Dirk Schlobinski, Lukas Schwintzer, Britta Qualmann, Michael M. Kessels

## Abstract

Local actin filament formation is indispensable for development of the dendritic arbor of neurons. We show that, surprisingly, the action of single actin filament-promoting factors was insufficient for powering dendritogenesis. Instead, this process required the actin nucleator Cobl and its only evolutionary distant ancestor Cobl-like acting interdependently. This coordination between Cobl-like and Cobl was achieved by physical linkage by syndapin I. Syndapin I formed nanodomains at convex plasma membrane areas at the base of protrusive structures and interacted with three motifs in Cobl-like, one of which was Ca^2+^/calmodulin-regulated. Consistently, syndapin I, Cobl-like’s newly identified N terminal calmodulin-binding site and the single Ca^2+^/calmodulin-responsive syndapin-binding motif all were critical for Cobl-like’s functions. In dendritic arbor development, local Ca^2+^/CaM-controlled actin dynamics thus relies on regulated and physically coordinated interactions of different F-actin formation-promoting factors and only together they have the power to bring about the sophisticated neuronal morphologies required for neuronal network formation.

## Introduction

The actin cytoskeleton is crucial for a huge variety of key processes in cell biology. Yet, only few factors were found that can promote the *de novo* formation of actin filaments (Chesarone & Goode 2009; Qualmann & Kessels, 2009). Thus, the initial idea that each of the discovered actin nucleators may be responsible for the formation of specific, perhaps tissue or cell-type-specific F-actin structures obviously had to be dismissed as too simple. Theoretically, the required functional diversity in actin filament formation despite a limited set of powerful effectors could be achieved by combinatory mechanisms specific for a given cell biological process. However, experimental evidence for such combinatory actions of actin nucleators still is very sparse. On top of that, which mechanisms may orchestrate these powerful effectors to bring about a certain cellular processes also remains a fundamental question in cell biology.

Neurons need to extend elaborate cellular protrusions - the single signal-sending axon and multiple signal-receiving, highly branched dendrites - to form neuronal networks. These very demanding and specialized cellular morphogenesis processes are driven by the actin cytoskeleton (Kessels et al., 2011). The formation of the dendritic arbor involves local Ca^2+^ and calmodulin (CaM) signals coinciding with transient F-actin formation by the evolutionary young actin nucleator Cobl (Cordon-bleu) (Ahuja et al., 2007) at dendritic branch induction sites (Hou et al., 2015). Ca^2+^/CaM regulates both Cobl’s loading with monomeric actin and its different modes of plasma membrane association (Hou et al., 2015). Cobl is furthermore regulated via arginine methylation by PRMT2 (Hou et al., 2018). All of these aspects were required for Cobl’s crucial role in dendritic arbor formation (Ahuja et al., 2007; Haag et al., 2012; Hou et al., 2015, Hou et al., 2018).

Interestingly, also Cobl’s evolutionary distant ancestor Cobl-like (COBLL1; Coblr1) was recently discovered to be important for Ca^2+^/CaM-controlled neuromorphogenesis (Izadi et al., 2018). While Cobl uses three Wiskott-Aldrich syndrome protein Homology 2 (WH2) domains to nucleate actin (Ahuja et al., 2007), Cobl-like employs a unique combinatory mechanism of G-actin binding by its single, C-terminal WH2 domain and Ca^2+^/CaM-promoted association with the actin-binding protein Abp1 (Kessels et al., 2000) to promote F-actin formation (Izadi et al., 2018). Cobl-like was also found to interact with Cyclin-dependent kinase 1 and to shape prostate cancer cells by not yet fully clear mechanisms (Takayama et al., 2018).

Here we show that Cobl and Cobl-like work at the same nascent dendritic branch sites in a strictly interdependent manner choreographed by physically bridging by the membrane-binding F-BAR protein syndapin I (Qualmann et al., 1999; Itoh et al., 2005; Dharmalingam et al., 2009; Schwintzer et al., 2011), which we identified to interact with Cobl-like and to specifically occur in nanoclusters at the convex membrane surfaces at the base of nascent membrane protrusions of developing neurons. The finding that one of the three syndapin binding sites of Cobl-like was regulated by Ca^2+^/CaM unveiled a further important mechanism of local control and coordination of actin dynamics in neuromorphogenesis.

Our work thereby provides insights into how two actin filament formation-promoting components - each critical for dendritic arbor formation - power actin-mediated dendritic branch initiation in a strictly coordinated manner and how this process can be directly linked to local membrane shaping to give rise to the complex morphologies required for proper neuronal network formation.

## Results

### Cobl-like and the actin nucleator Cobl work at the same dendritic branching sites and largely phenocopy each other in their critical role in dendritic arborization

The actin nucleator Cobl and its evolutionary ancestor protein Cobl-like are molecularly quite distinct (**Figure 1–figure supplement 1**), however, both critical for dendritic arbor formation (Ahuja et al., 2007; Izadi et al., 2018). Side-by-side loss-of-function analysis of both factors in developing primary hippocampal neurons using IMARIS software-based evaluations for detailed analyses of the elaborate morphology of such cells (Izadi et al., 2018) revealed surprisingly similar phenotypes (**Figure 1A-G**). Dendritic branch and terminal point numbers as well as total dendritic length all were severely affected by lack of Cobl-like (**Figure 1A-G**). Corresponding Cobl loss-of-function showed that, while dendritic growth processes seemed largely unaffected by Cobl deficiency, also Cobl deficiency led to a significant reduction of terminal points and in particular to severe loss of dendritic branch points. With −35%, these defects were about as strong as those caused by Cobl-like RNAi (**Figure 1D-F**). Cobl RNAi mostly affected Sholl intersections in proximal areas. Cobl-like RNAi led to reduced Sholl intersections throughout the dendritic arbor (**Figure 1G**).

**Figure 1.**
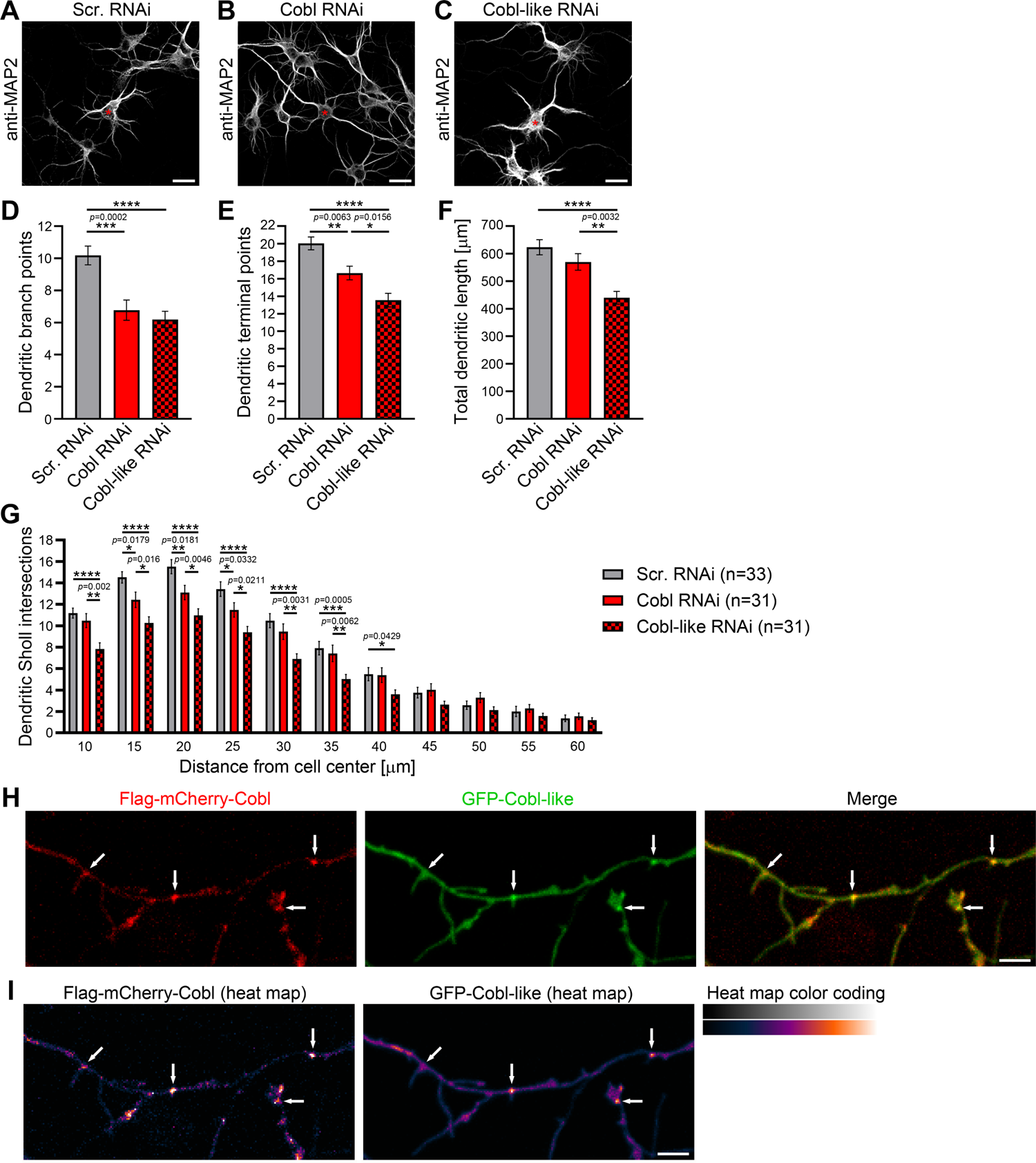
Cobl-like and the actin nucleator Cobl work at the same dendritic branching sites and largely phenocopy each other in their critical role in dendritic arborization. (**A-C**) Maximum intensity projections (MIPs) of anti-MAP2 immunostained developing primary hippocampal neurons transfected as indicated at DIV4 and fixed at DIV5.5. Asterisks, transfected neurons. Bars, 20 µm. (**D-G**) Quantitative comparative Cobl-like and Cobl loss-of-function analyses of indicated dendritic parameters. (**H, I**) MIPs of GFP-Cobl-like and Flag-mCherry-Cobl in dendrites of developing hippocampal neurons (DIV6) in standard colors (**H**) and as heat map representation (**I**), respectively. Arrows, examples of putative, nascent dendritic branch induction sites with accumulations of both Cobl and Cobl-like. Bar, 5 µm. Data, mean±SEM. 1-way ANOVA+Tukey (**D-F**) and 2-way ANOVA+Bonferroni (**G**).

The phenotypical comparison of Cobl and Cobl-like unveiled that both cytoskeletal effectors have somewhat similar functions in dendritic arborization. Colocalization studies showed that Flag-mCherry-Cobl and GFP-Cobl-like did not show any obvious spatial segregation (neither in proximal nor in peripheral dendritic arbor of developing neurons) but largely colocalized. Dendritic accumulations of Cobl usually showed corresponding albeit less pronounced accumulations of Cobl-like (**Figure 1H,I**; arrows). This suggested that Cobl and Cobl-like are not responsible for distinct branching sites but work at the same sites.

### Cobl-like functions in dendritic arborization strictly depend on Cobl and likewise Cobl functions depend on Cobl-like

Two powerful molecular machines for actin filament formation at the same place may either reflect function redundancy or parallel action to drive cellular processes effectively in response to (putatively different) signaling cues or may even reflect interlinked functions. Functional redundancy seemed unlikely, because both individual loss-of-function phenotypes were severe (Cobl, −33%; Cobl-like, −39%) (**Figure 1D**). Thus, we focused on the remaining hypotheses.

Interestingly, Cobl-like-driven dendritic arborization was completely impaired by the lack of Cobl (**Figure 2A-G**). Dendritic branch point numbers, terminal point numbers and the total dendritic arbor length of neurons cotransfected with GFP-Cobl-like and Cobl RNAi all were highly statistically significantly below those of neurons cotransfected with Cobl-like and scrambled RNAi. The suppression of the Cobl-like gain-of-function effects by Cobl RNAi was so strong that under the chosen condition (34 h expression; GFP coexpression) all three parameters were indistinguishable from control levels and therefore permitted the conclusion that the effects reflected a full suppression of Cobl-like’s functions in dendritic branching in the absence of Cobl (**Figure 2E-G**).

**Figure 2.**
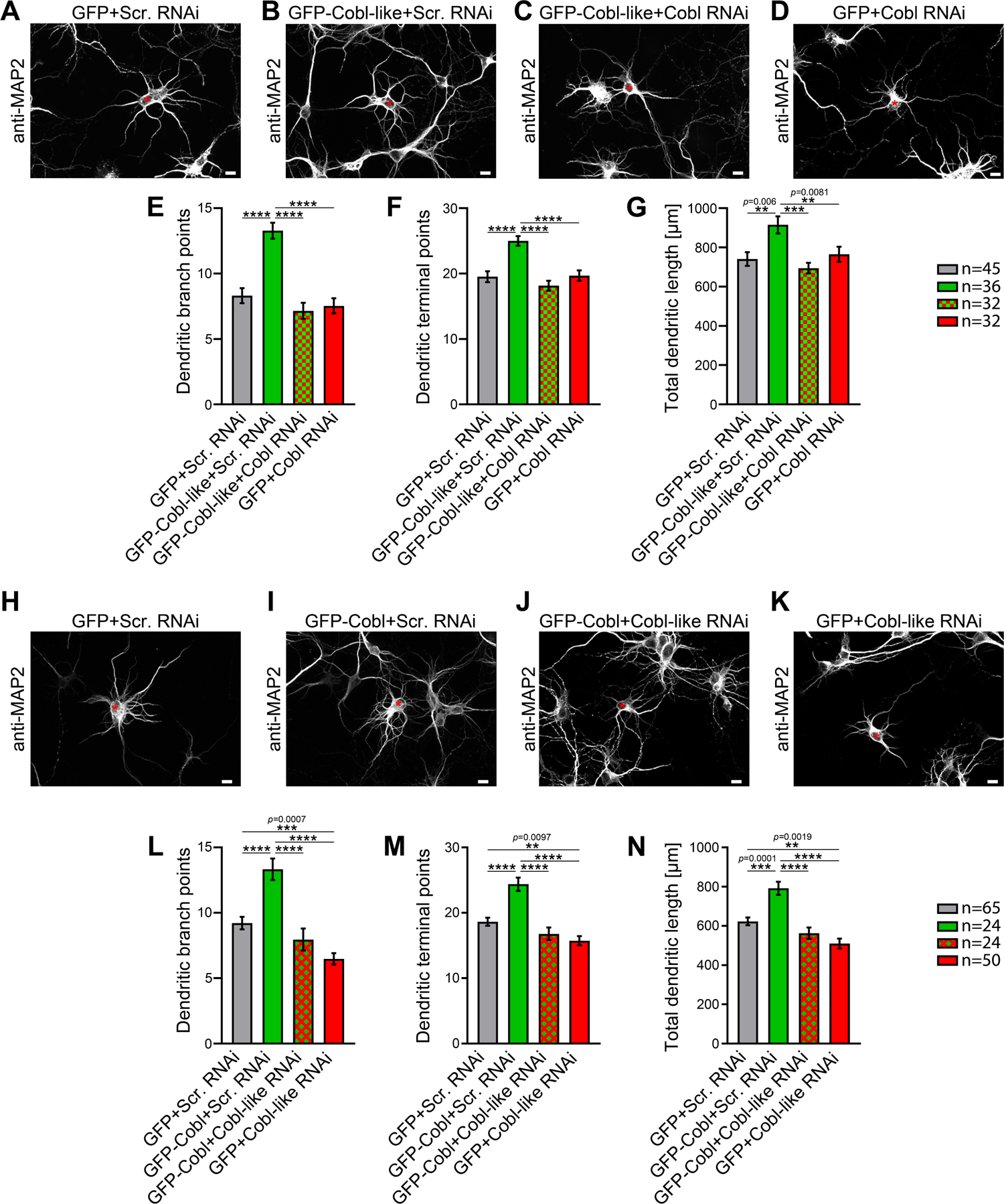
Functional interdependence of Cobl-like and Cobl in dendritic arbor formation. (**A-D**) MIPs of neurons showing the suppression of the Cobl-like gain-of-function phenotype (**B**; GFP-Cobl-like+Scr. RNAi) by mCherryF-reported RNAi plasmids directed against Cobl (**C**) in comparison to control neurons (**A**; GFP+Scr. RNAi) and Cobl RNAi neurons (**D**; GFP+Cobl RNAi). (**E-G**) Quantitative determinations of indicated dendritic arborization parameters unveiling a full suppression of all Cobl-like functions in dendritic arbor formation by a lack of Cobl. (**H-N**) Related images (**H-K**) and quantitative data (**L-N**) of experiments revealing a functional dependence of Cobl on Cobl-like. Asterisks, transfected neurons. Bars, 10 µm. Data, mean±SEM. 1-way ANOVA+Tukey (**E-G** and **L-N**).

To our surprise, likewise, also Cobl-promoted dendritic arbor formation turned out to be massively affected by absence of Cobl-like (**Figure 2H-K**). The dendritic parameters of developing neurons expressing Cobl and Cobl-like RNAi did not show any Cobl gain-of-function phenotype but were statistically not significantly different from those of control or Cobl-like RNAi (**Figure 2L-N**). Thus, Cobl functions in dendritic arbor formation clearly depended on Cobl-like.

Taken together, Cobl and Cobl-like both are cellular factors promoting actin filament formation and significantly differ in their properties, yet, in dendritic branch formation, they do not work independently but surprisingly strictly depend on each other.

### Cobl-like associates with syndapins

The surprising functional interdependence of Cobl and Cobl-like in dendritic arbor formation raised the question how this may be organized mechanistically with two proteins that seem to employ quite different molecular mechanisms (Ahuja et al., 2007; Izadi et al., 2018).

Using a variety of different methods, we failed to observe any obvious interactions of Cobl and Cobl-like (our unpublished efforts; also see below). Thus, the crosstalk of Cobl-like and Cobl had to be less direct and more sophisticated.

Cobl was demonstrated to use complexly choreographed membrane binding mechanisms involving its direct binding partner syndapin I (Schwintzer et al., 2011; Hou et al., 2015). Syndapins can self-associate (Kessels and Qualmann, 2006) and could therefore theoretically link Cobl and Cobl-like physically. As a prerequisite, syndapin I would have to associate with Cobl-like. Indeed, GFP-Cobl-like specifically coprecipitated with immobilized syndapin I SH3 domain. The interaction was mediated by N terminal proline-rich regions of Cobl-like (**Figure 3A**) and was conserved among syndapin I, syndapin II and syndapin III (**Figure 3B**).

**Figure 3.**
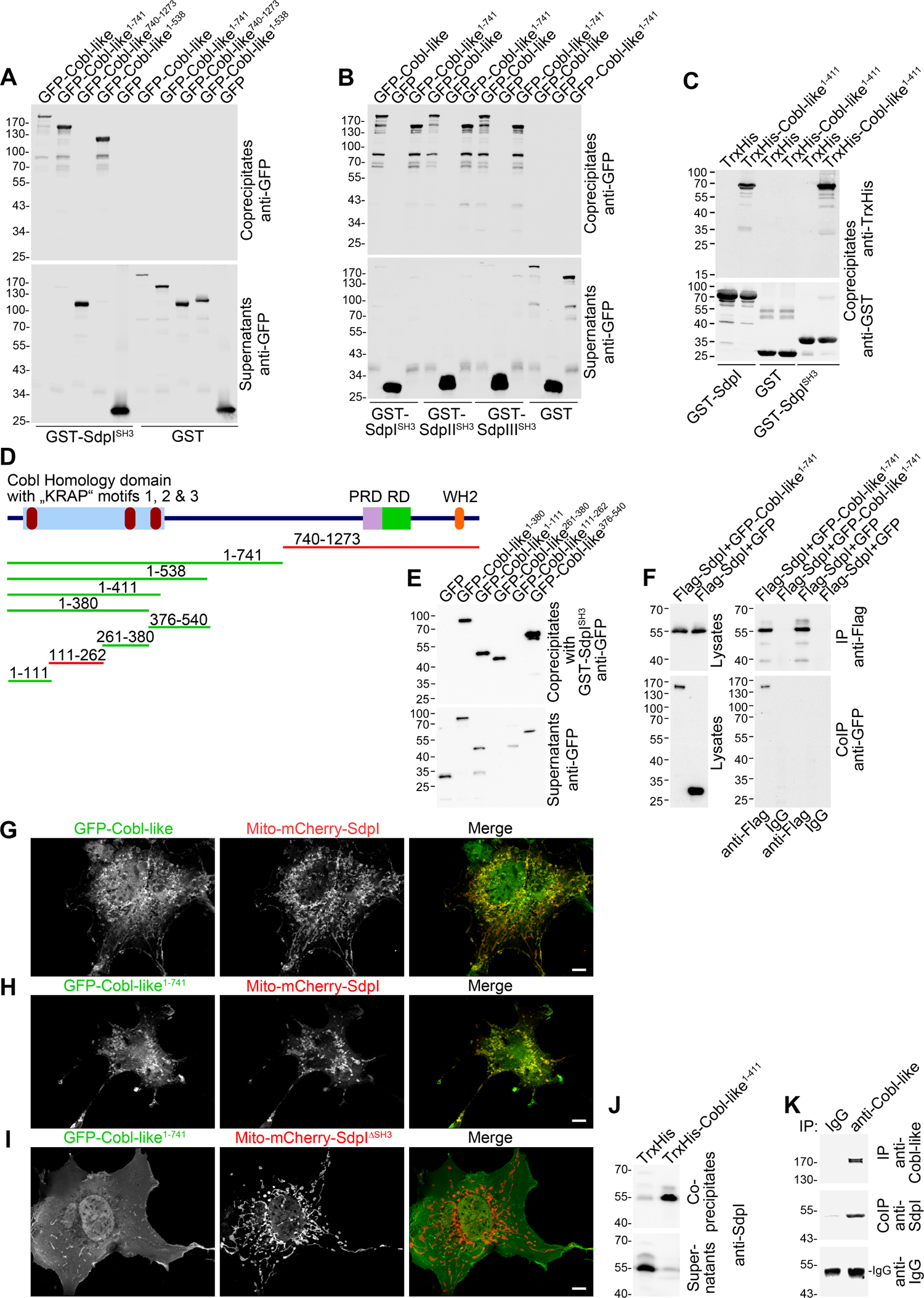
Cobl-like associates with syndapins. (**A**) Coprecipitation analyses of GFP-tagged Cobl-like and deletion mutants thereof with immobilized syndapin I SH3 domain (GST-SdpI^SH3^). (**B**) Related coprecipitation studies with the SH3 domains of syndapin I, syndapin II (SdpII^SH3^) and syndapin III (SdpIII^SH3^), respectively. (**C**) Reconstitution of the association of TrxHis-Cobl-like^1-411^ with purified GST-syndapin I and GST-syndapin I SH3 domain but not with GST. (**D**) Scheme of Cobl-like with its domains (PRD, proline-rich domain; RD, Abp1-binding repeat domain; WH2, WH2 domain) and deletion mutants used (red, not binding syndapins; green, binding). Red in the Cobl Homology domain, “KRAP” motifs (not drawn to scale). (**E**) Coprecipitation assays with Cobl-like deletion mutants mapping Cobl-like’s syndapin binding sites. (**F**) Coimmunoprecipitations unveiling a specific association of GFP-Cobl-like^1-741^ with Flag-syndapin I. (**G-I**) Reconstitution and visualization of Cobl-like/syndapin I complexes in COS-7 cells using mitochondrially targeted syndapin I (**G,H**) as well as a mutant lacking the SH3 domain (Mito-mCherry-SdpI^ΔSH3^) (**I**) with GFP-Cobl-like (**G**) and GFP-Cobl-like^1-741^ (**H,I**). Bars, 10 µm. (**J**) Coprecipitation of endogenous syndapin I from mouse brain lysates by TrxHis-Cobl-like^1-411^. (**K**) Endogenous coimmunoprecipitation of Cobl-like and syndapin I from mouse brain lysates.

*In vitro* reconstitutions with purified components proved that syndapin I/Cobl-like interactions were direct (**Figure 3C**) and were furthermore based on classical SH3 domain/PxxP motif interactions, as proven by using a P434L-mutated SH3 domain (**Figure 3–figure supplement 1A**).

Cobl-like deletion mutants (**Figure 3D,E**) showed that specifically three regions in Cobl-like’s Cobl Homology domain (Cobl-like^1-111^, Cobl-like^261-380^ and Cobl-like^376-540^) contained syndapin I interfaces (**Figure 3E, Figure 3–figure supplement 1B**). Each of them has a single “KRAP” motif (**Figure 1–figure supplement 1**).

Specific coimmunoprecipitation of GFP-Cobl-like^1-741^ with Flag-tagged syndapin I demonstrated that the identified interaction can also occur *in vivo* (**Figure 3F**). GFP-Cobl-like^1-741^ also specifically coimmunoprecipitated with Flag-syndapin II-s and Flag-syndapin III (**Figure 3–figure supplement 1C,D**), i.e. with syndapin family members showing a wider expression than syndapin I (Kessels and Qualmann, 2004).

It was furthermore possible to directly visualize Cobl-like/syndapin I complex formation in intact cells by demonstrating specific recruitments of GFP-Cobl-like and GFP-Cobl-like^1-741^ to mitochondria decorated with Mito-mCherry-syndapin I (**Figure 3G,H; Figure 3–figure supplement 1E)**. This firmly excluded postsolubilization artefacts, which theoretically could compromise biochemical studies. Deletion of the syndapin I SH3 domain (Mito-mCherry-SdpI^ΔSH3^) disrupted complex formations with both GFP-Cobl-like^1-741^ (**Figure 3I**) and GFP-Cobl-like full-length (**Figure 3–figure supplement 1F**).

Cobl-like/syndapin interactions also are of relevance in the brain, as immobilized, recombinant TrxHis-tagged Cobl-like specifically precipitated endogenous syndapin I from mouse brain lysates (**Figure 3J**). Furthermore, endogenous Cobl-like/syndapin I complexes *in vivo* were demonstrated by coimmunoprecipitation analyses from mouse brain lysates (**Figure 3K**).

### Syndapin I is crucial for Cobl-like’s ability to promote dendritic arbor extension and branching

We next addressed whether the identified Cobl-like interaction partner syndapin I would indeed be critical for Cobl-like’s functions. GFP-Cobl-like massively promoted dendritic arborization already after very short times (Izadi et al., 2018). Strikingly, all Cobl-like gain-of-function phenotypes in developing primary hippocampal neurons were completely suppressed upon syndapin I RNAi (**Figure 4C**). Cobl-like-overexpressing neurons cotransfected with syndapin I RNAi showed dendritic branch points, dendritic terminal points and an overall length of the dendritic arbor that were highly statistically significantly different from Cobl-like overexpression neurons and indistinguishable from those of control cells. The syndapin I RNAi-mediated suppression of Cobl-like functions occurred in all dendritic arbor parts affected by Cobl-like gain-of-function (**Figure 4D-G**). Cobl-like’s functions in dendritic arbor formation thus are fully dependent on the availability of its direct interaction partner syndapin I.

**Figure 4.**
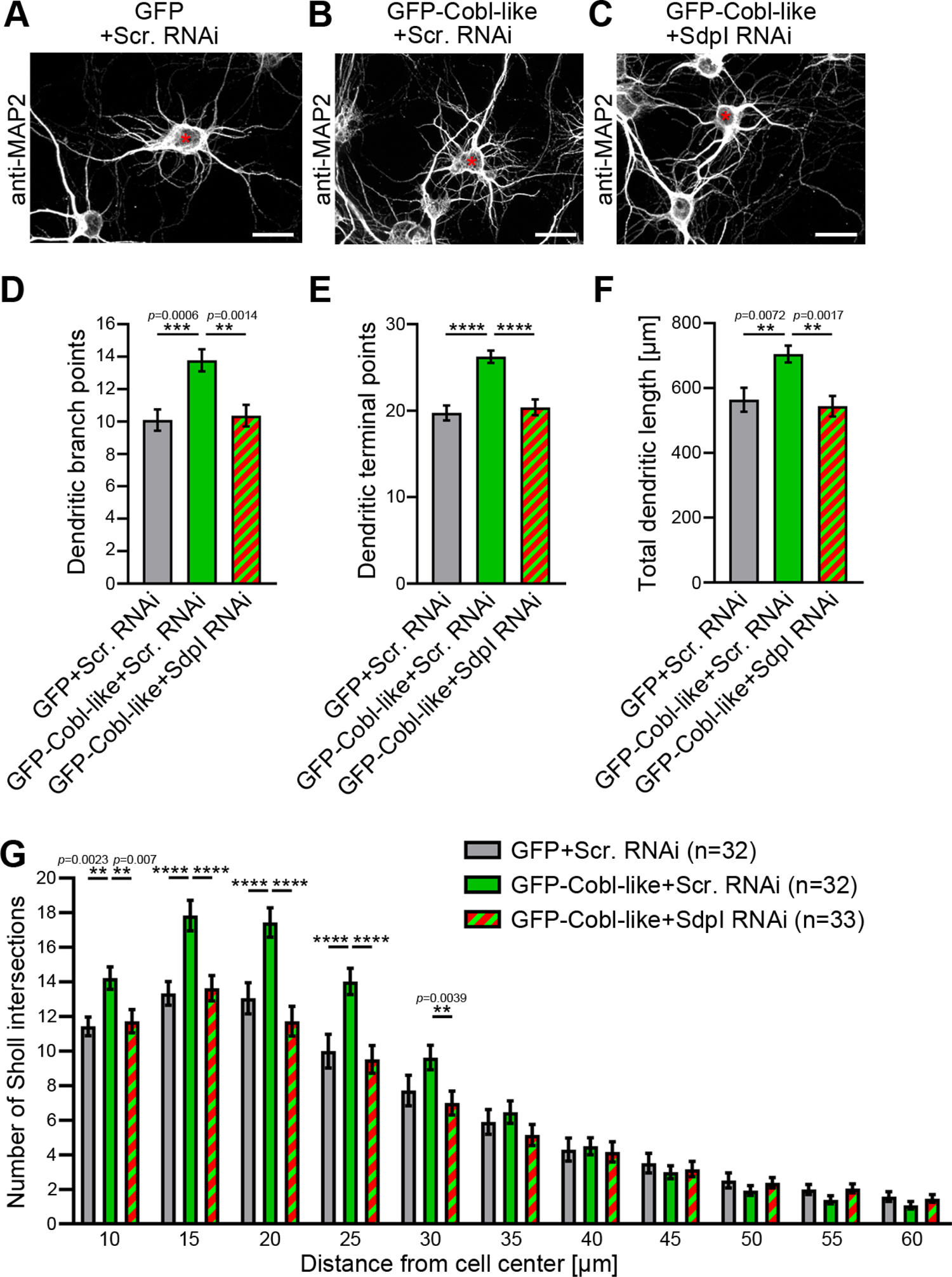
Cobl-like functions in dendritic arbor formation rely on syndapin I. (**A-C**) MIPs of DIV5.5 neurons transfected as indicated. Asterisks, transfected neurons. Bars, 20 µm. (**D-G**) Quantitative determinations of key dendritic arborization aspects promoted by Cobl-like for their dependence on syndapin I. Data, mean±SEM. 1-way ANOVA+Tukey (**D-F**) and 2-way ANOVA+Bonferroni (**G**).

### Syndapins physically interconnect Cobl-like with Cobl

In order to unravel molecular mechanisms underlying the strict functional interdependence of Cobl and Cobl-like in dendritic arborization, we asked whether syndapin I may indeed be able to directly bridge the two actin cytoskeletal effectors. To exclude putative indirect interactions via actin, we used immobilized GST-Cobl-like^1-411^, which comprises the syndapin binding sites (**Figure 3**). Cobl-like^1-411^ indeed formed specific protein complexes with GFP-Cobl^1-713^ when syndapin I was present (**Figure 5A**). No GFP-Cobl^1-713^ was precipitated when syndapin I was omitted. Thus, direct interactions between Cobl-like and Cobl did not occur but complex formation required syndapin I acting as a bridge (**Figure 5A**). Likewise, also syndapin III mediated complex formation of Cobl-like with Cobl (**Figure 5B**).

**Figure 5.**
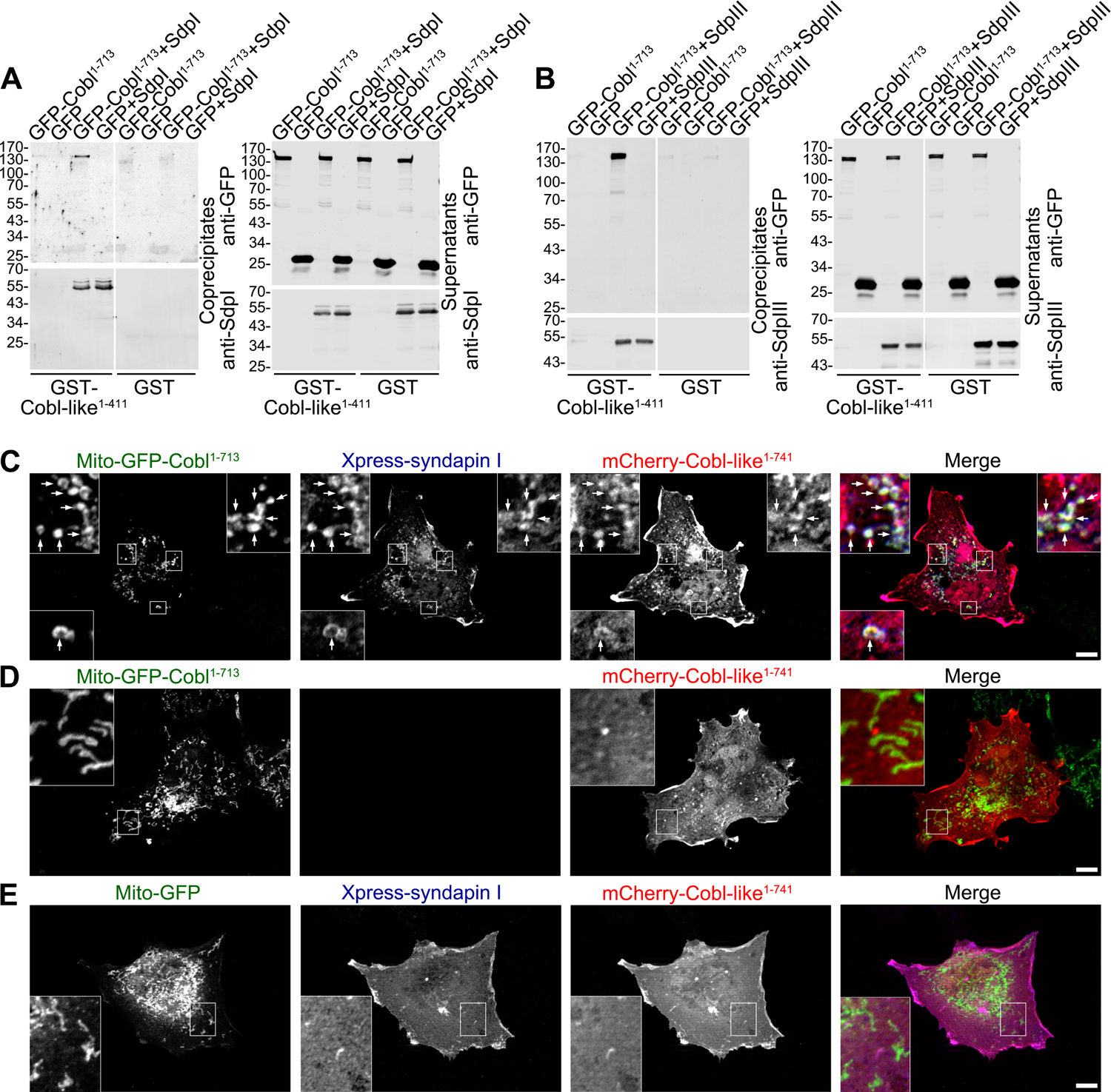
Cobl-like is physically linked to Cobl via syndapin I acting as a bridging component. (**A,B**) Coprecipitation analyses unveiling specific and syndapin-dependent formation of complexes composed of immobilized GST-Cobl-like, syndapin I (**A**) and syndapin III (**B**), respectively, as well as GFP-Cobl^1-713^. White lines indicate omitted blot lanes. (**C-E**) Reconstitution and visualization of Cobl-like/syndapin I/Cobl protein complexes in COS-7 cells. Mito–GFP-Cobl^1-713^ (**C,D**) but not Mito-GFP (**E**) recruited mCherry-Cobl-like^1-741^ in the presence of Xpress-syndapin I (**C**) but not in its absence (**D**). Boxes in **C-E,** areas presented as magnified insets (**C,D**, 4fold; **E**, 3fold). Arrows, examples of colocalization of all three channels. Bars, 10 µm.

Demonstrating that complexes of all three components are also formed at membranes and in intact cells Mito-GFP-Cobl^1-713^ successfully recruiting syndapin I to mitochondrial membranes (**Figure 5–figure supplement 1**) also recruited Cobl-like^1-741^ (**Figure 5C**). The visualized complex formations were specific and mediated by syndapin I acting as bridging component between Cobl and Cobl-like, as omitting syndapin I did not lead to any Cobl-like^1-741^ mitochondrial presence and also Mito-GFP did not lead to any syndapin/Cobl-like colocalization at mitochondria (**Figure 5D,E**).

### Syndapin I and Cobl-like colocalize at sites of dendritic branch induction

In line with the BAR domain hypothesis (Peter et al., 2004; Qualmann et al., 2011; Daumke et al., 2014; Kessels and Qualmann, 2015; Carman and Dominguez, 2018) syndapin I may sense/induce certain membrane topologies and thereby provide spatial cues for Cobl and Cobl-like functions. Thus, three syndapin I-related aspects needed to be experimentally addressed: i) Where and when do Cobl-like and syndapin I occur together in developing neurons? ii) Would a given syndapin I localization indeed reflect specifically membrane-associated syndapin I? iii) Would putative accumulations of membrane-associated syndapin I then really correspond to convex membrane topologies?

Dual time-lapse imaging of GFP-Cobl-like and syndapin I-mRubyRFP in developing primary hippocampal neurons showed that syndapin I accumulated in defined spots along dendrites coinciding with subsequent branch induction events. Such accumulations occurred as early as 1 min prior to detectable dendritic branch protrusion, were spatially restricted to very small areas (diameters, ∼250-1200 nm) and were spatially and temporally colocalized with Cobl-like at branch initiation sites (**Figure 6A,B**).

**Figure 6.**
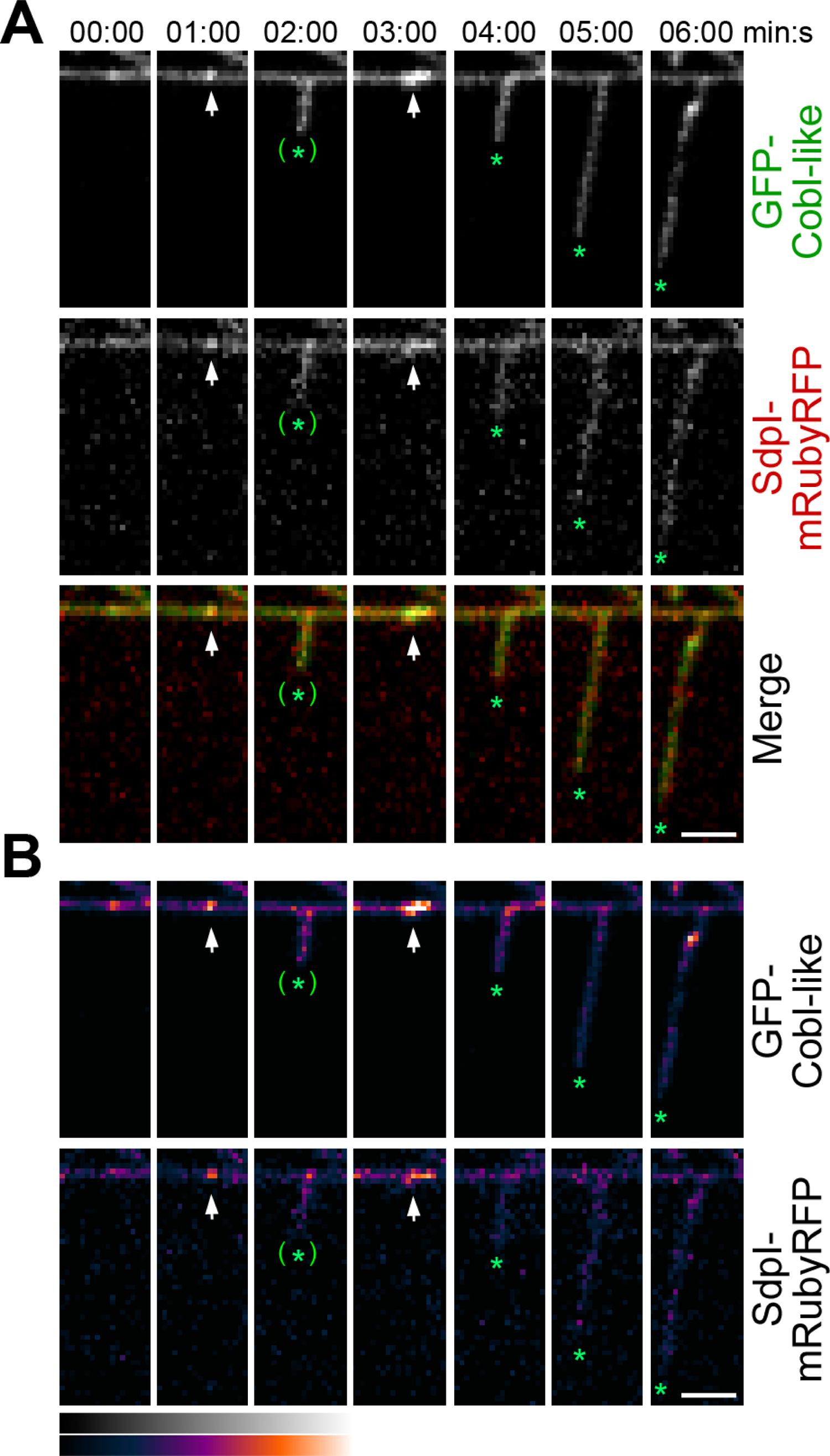
Cobl-like and syndapin I coincide at nascent dendritic branch sites. **(A)** MIPs of individual frames of a 3D-time-lapse recording of a dendrite segment of a DIV7 rat hippocampal neuron coexpressing GFP-Cobl-like and syndapin I (SdpI)-mRubyRFP. Arrows, GFP-Cobl-like and syndapin I enrichments prior to protrusion initiation from these dendritic sites; *, tips of growing dendritic protrusions; (*), abandoned protrusions. (**B**) Heat map representations. Bars, 2.5µm.

Afterwards, the accumulations of both proteins at the base of newly formed protrusions faded. This suggested a highly mobile subpool of syndapin I and Cobl-like in the dendritic arbor. Sites with repetitive dendritic protrusion attempts showed accumulations of both syndapin I and Cobl-like prior to the first as well as prior to the second protrusion initiation (**Figure 6A,B**).

### Membrane-bound syndapin I occurs preferentially at protrusive membrane topologies in developing neurons and forms nanoclusters at such sites

3D-time-lapse studies do not resolve whether the observed syndapin I accumulations at nascent branch sites represent membrane-associated syndapin I or a cytosolic subpool e.g. associated with putative cytoskeletal components at such sites. Immunogold labeling of freeze-fractured plasma membranes is a technique that *per se* exclusively focusses on membrane-integrated proteins, provides membrane topology information, and can be applied to neuronal networks (e.g. see Tanaka et al., 2005; Holderith et al., 2012; Schneider et al., 2014; Nakamura et al., 2015). We have recently shown that plasma membranes of still developing neurons can in principle be freeze-fractured and immunolabelled, too (Wolf et al., 2019).

Whereas either Cobl nor Cobl-like seemed preserved by the procedure (our unpublished data), immunogold labeling of syndapin I, which can insert hydrophobic wedges into one membrane leaflet (Wang et al., 2009), was successfully obtained (**Figure 7A-B**). In principle, anti-syndapin I immunogold labeling was seen at both cylindrical and protrusive membrane topologies. However, even at the conditions of saturated labeling applied, cylindrical membrane surfaces merely showed sparse anti-syndapin I immunogold labeling and were mostly decorated with single gold particles or by pairs of labels. In contrast, at protrusive sites, the labeling density was about three times as high as at cylindrical surfaces (**Figure 7A,B**).

**Figure 7.**
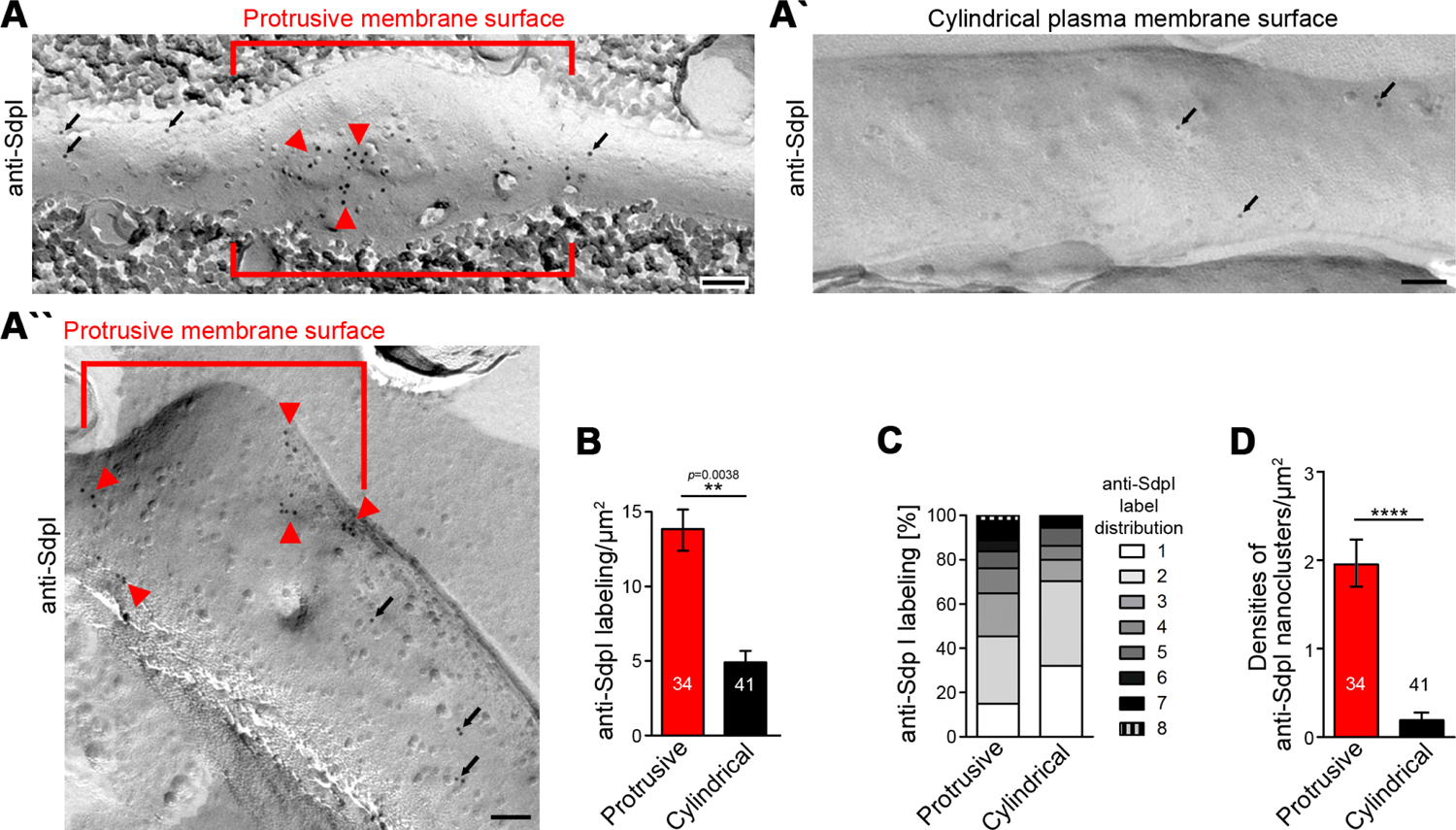
Syndapin I nanoclusters are enriched at sites of dendritic protrusion. (**A,A’,A’’**) TEM images of anti-syndapin I immunogold-labeled freeze-fracture replica of developing neurons (DIV7). Red lines highlight membrane topologies protruding from regular cylindrical topology. Arrowheads, abundant and clustered anti-syndapin I immunogold labeling (10 nm) at protrusive sites. Arrows, sparse and rarely clustered anti-syndapin immunogold labeling at regular, cylindrical membrane structures. Bars, 200 nm. (**B**) Quantitative evaluations of anti-syndapin I labeling densities at protrusive and cylindrical membrane topologies. (**C**) Quantitative analysis of the relative abundance of differently clustered syndapin I labels (ROIs, 35 nm radius). In total, 335 (protrusive) and130 (cylindrical) labels were evaluated. (**D**) Quantitative analysis of the density of anti-syndapin I nanoclusters (≥3 anti-syndapin I immunogold labels/ROI) at regular cylindrical membrane surfaces and at those with protrusive topology. Data **B,D**, mean±SEM. 1-way ANOVA (**B**); two-tailed Student’s t-test (**D**).

Protrusive sites also showed a statistically highly significant enrichment of syndapin I nanoclusters (≥3 anti-syndapin I labels in circular ROIs of 35 nm radius) (**Figure 7B-D**). Interestingly, syndapin I was usually not localized to the tip of the protrusion but preferentially occurred at membrane topology transition zones at the protrusion base (**Figure 7A’’**). The accumulation of syndapin I clusters at such sites was in line with a promotion of membrane curvature induction and/or with a stabilization of the complex membrane topologies found at such sites by syndapin I.

### Cobl-like’s N terminus is a target for the Ca^2+^ sensor CaM and Ca^2+^ signals increase Cobl-like’s associations with syndapin I

The formation of neuronal networks involves local Ca^2+^ and CaM signals, which coincide with transient F-actin formation at sites of dendritic branch induction (Hou et al., 2015). Cobl-like was identified as Ca^2+^/CaM target. Yet, this CaM association occurred in the C terminal part of Cobl-like and regulated Cobl-like’s association with the F-actin-binding protein Abp1 (Izadi et al., 2018).

Interestingly, also GFP-Cobl-like^1-411^ showed Ca^2+^-dependent CaM binding, whereas middle parts, such as Cobl-like^376-540^ and Cobl-like^537-740^, did not (**Figure 8A,B**). Surprisingly, further analyses demonstrated that the central parts of the Cobl Homology domain of Cobl-like, i.e. Cobl-like^111-262^ and Cobl-like^182-272^ also both did not show any Ca^2+^-dependent CaM binding (**Figure 8A,B**), although the central Cobl Homology domain corresponds to the CaM-binding area in Cobl (Hou et al., 2015) and represents an area of at least moderately higher sequence conservation between Cobl and Cobl-like (33% identity; **Figure 1–figure supplement 1**). Instead, it was the most N-terminal part of Cobl-like represented by Cobl-like^1-111^ and Cobl-like^1-58^ that was targeted by CaM (**Figure 8A,B**).

**Figure 8.**
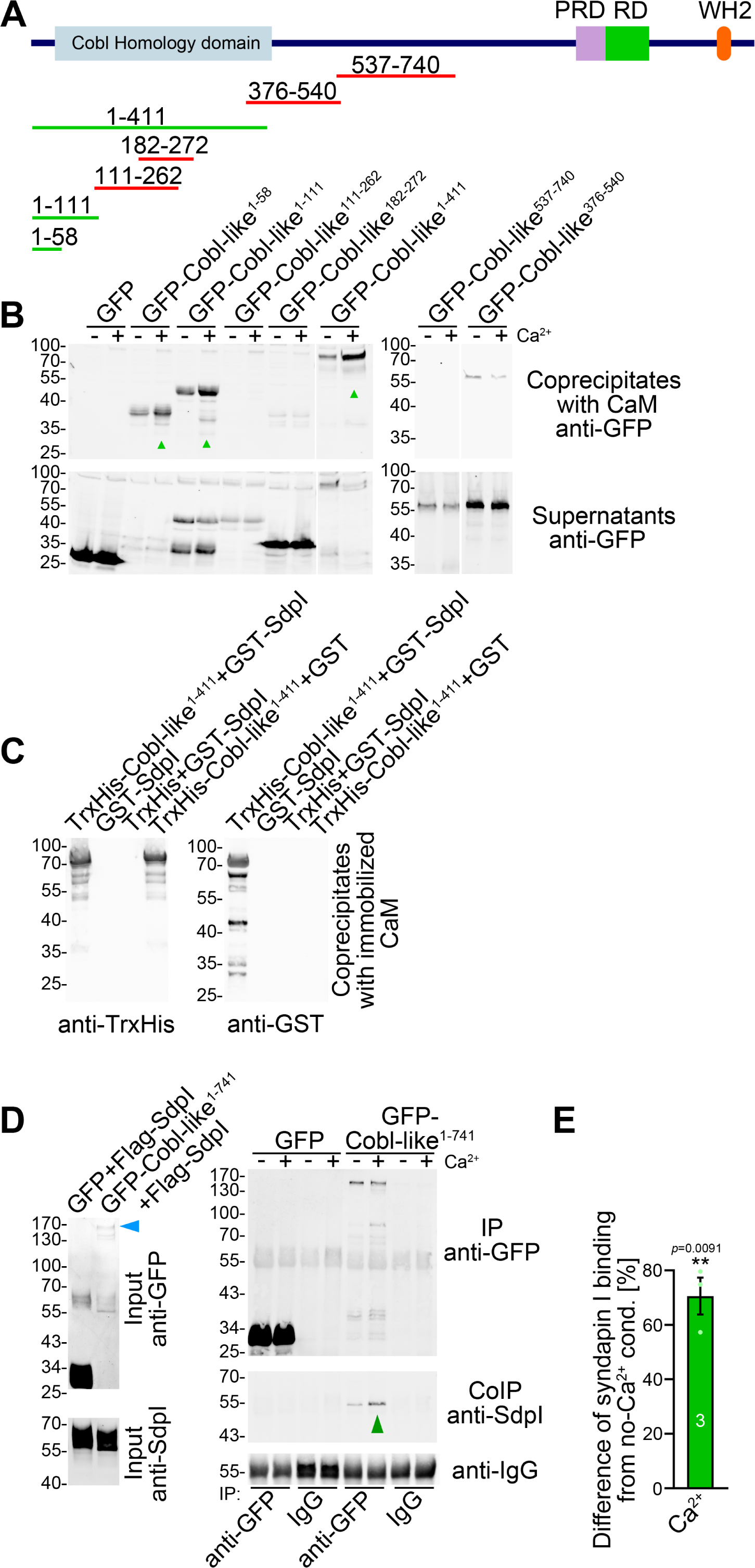
Ca^2+^/CaM associates with the N terminus of Cobl-like and positively regulates Cobl-like’s syndapin I association. (**A**) Scheme of Cobl-like and deletion mutants used for CaM binding studies (**B**) (red, no Ca^2+^-dependent binding; green, Ca^2+^-dependent binding). (**B**) Coprecipitations with immobilized CaM in presence (500 µM) and absence of Ca^2+^ and different Cobl-like deletion mutants. Green arrowheads, increased CaM interactions in the presence of Ca^2+^. White lines, lanes omitted from blots. (**C**) Coprecipitation analyses with immobilized CaM and purified TrxHis-Cobl-like^1-411^ and GST-syndapin I (GST-SdpI) showing direct and simultaneous interactions of Cobl-like^1-411^ with both CaM and syndapin I. (**D,E**) Quantitative coimmunoprecipitation analyses demonstrating that Ca^2+^/CaM signaling leads to increased syndapin I coimmunoprecipitation with Cobl-like^1-741^. Blue arrowhead, position of the only faintly detected GFP-Cobl-like^1-741^ in the lysates (**D**). Green arrowhead, increase of coimmunoprecipitated syndapin I (**D**). (**E**) Anti-syndapin I signal per immunoprecipitated Cobl-like (expressed as change from conditions without Ca^2+^). Data, bar/dot plot overlays with mean±SEM. Unpaired student’s t-test.

Coprecipitation experiments with purified recombinant proteins confirmed that Ca^2+^/CaM and syndapin I can bind Cobl-like^1-411^ simultaneously (**Figure 8C**). We hypothesized that the discovered Ca^2+^/CaM binding to the Cobl-like N terminus may play a role in regulating the syndapin binding of the neighbored Cobl Homology domain. Quantitative syndapin I coimmunoprecipitation experiments with Cobl-like^1-741^ demonstrated an improved complex formation of Cobl-like with syndapin I when Ca^2+^ was added. With an increase of ∼70%, syndapin I binding to Cobl-like^1-741^ turned out to be massively promoted by Ca^2+^ (**Figure 8D,E**). Thus, the identified N terminal complex formation with syndapin I is Ca^2+^/CaM-regulated.

### Cobl-like’s N terminal CaM binding site regulating syndapin I association levels is crucial for dendritic arbor formation

The N terminal region of Cobl-like (**Figure 1–figure supplement 1**) indeed contains putative CaM binding motifs. Coprecipitation analyses clearly showed that, in contrast to GFP-Cobl-like^1-741^, a corresponding deletion mutant (GFP-Cobl-like^1-741ΔCaM NT^; GFP-Cobl-like^1-741Δ11-45^) did not show any Ca^2+^-dependent CaM binding (**Figure 9A**).

**Figure 9.**
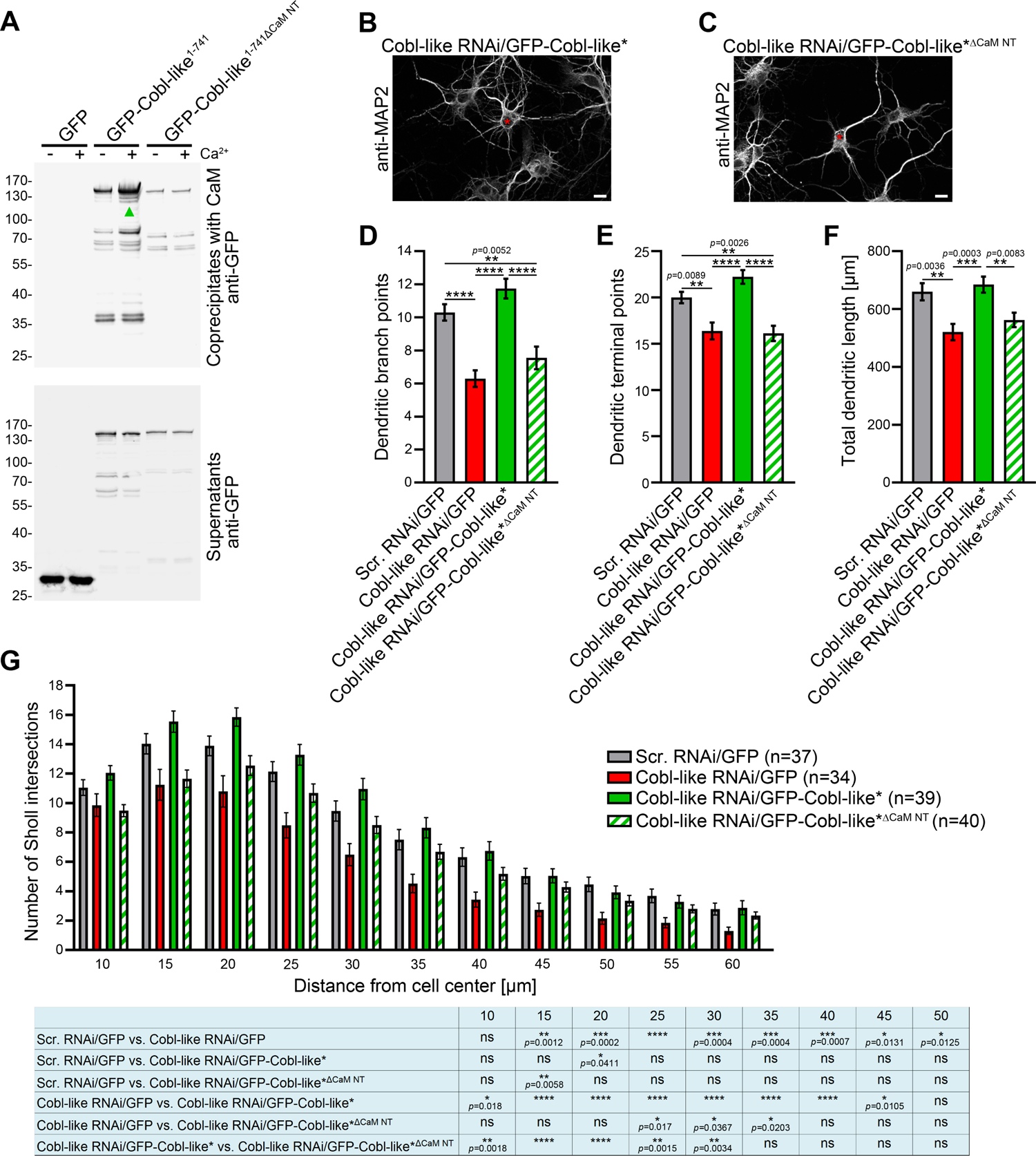
The N terminal CaM binding site of Cobl-like is indispensable for all critical functions of Cobl-like in dendritic arbor formation. (**A**) Coprecipitation analyses of Cobl-like^1-741^, Cobl-like^1-741ΔCaM NT^ (11-45) and GFP with immobilized CaM in Ca^2+^ presence and absence. Arrowhead, increased CaM interaction of Cobl-like^1-741^ upon Ca^2+^ (disrupted in Cobl-like^1-741ΔCaM NT^). (**B-C**) Functional analyses in primary hippocampal neurons unveiling that an RNAi-insensitive (*) Cobl-like mutant lacking the N terminal CaM binding site (GFP-Cobl-like*^ΔCaM NT^) failed to rescue the Cobl-like loss-of-function phenotypes. Red asterisks, transfected neurons (transfection, DIV4; analyses, DIV5.5). Bars, 10 µm. (**D-G**) Quantitative evaluations of indicated dendritic parameters. Data, mean±SEM. 1-way ANOVA+Tukey (**D-F**); 2-way ANOVA+Bonferroni (**G**).

Strikingly, a RNAi-resistant (*) Cobl-like mutant solely lacking the N terminal CaM binding site (GFP-Cobl-like*^ΔCaM NT^) failed to rescue the Cobl-like loss-of-function phenotypes in dendritic arborization (**Figure 9B,C**). Quantitative analyses unveiled that reexpression of GFP-Cobl-like*^ΔCaM NT^ instead of resupplying the neurons with RNAi-insensitive wild-type Cobl-like*, which rescued all Cobl-like deficiency phenotypes, led to defects in dendritic branch point numbers, terminal point numbers and total dendritic length. These defects were as severe as those caused by Cobl-like RNAi without rescue attempt (**Figure 9E-F**).

Also Sholl analyses confirmed that GFP-Cobl-like*^ΔCaM NT^ showed a significant lack of performance in all proximal and central parts of the dendritic arbor when compared to Cobl-like RNAi/GFP-Cobl-like* (statistically significantly different at all Sholl intersections up to 30 µm) (**Figure 9G**). The identified N terminal CaM binding site of Cobl-like regulating the syndapin I interactions thus was absolutely indispensable for Cobl-like’s functions in dendritic arbor formation.

### Ca^2+^/CaM signaling exclusively promotes the syndapin I association with the first of the three “KRAP” motifs

The critical N terminal CaM binding site was adjacent to the most N terminal of the three syndapin binding areas. As a prerequisite for further analyses uncovering the regulatory mechanism, we next confirmed that the interactions with syndapins were indeed solely mediated by the “KRAP” motif-containing regions. Both Cobl-like^1-741ΔKRAP^ and Cobl-like^ΔKRAP^ indeed were not able to interact with the syndapin I SH3 domain, as shown by coprecipitation studies and by reconstitutions of complex formations with syndapin I *in vivo* (**Figure 10A,B**; **Figure 10–figure supplement 1**).

**Figure 10.**
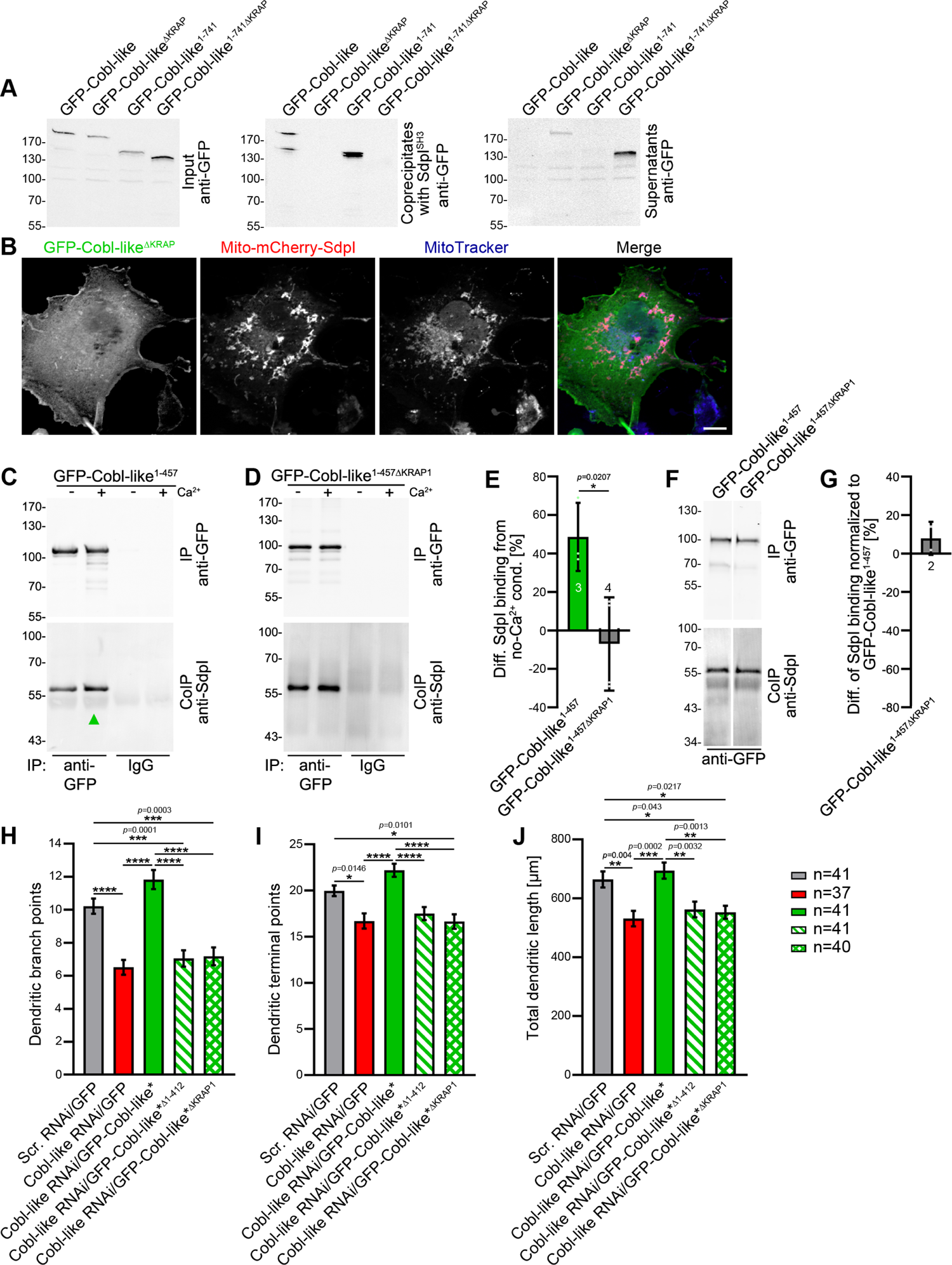
Ca^2+^/CaM signaling exclusively promotes the syndapin I association with the first of the three “KRAP” motifs and this single Ca^2+^/CaM-regulated motif is crucial for Cobl-like’s functions. (**A**) Coprecipitations with immobilized syndapin I SH3 domain (SdpI^SH3^) and Cobl-like and ΔKRAP mutants thereof. (**B**) MIP of a MitoTracker-stained COS-7 cell transfected with Mito-mCherry-syndapin I and GFP-Cobl-like^ΔKRAP^ (colocalization of only red and blue channels, purple in merge). Bar, 10 µm. (**C-G**) Quantitative coimmunoprecipitation analyses with GFP-Cobl-like^1-457^ (**C**) in comparison to a corresponding mutant solely lacking the first “KRAP” motif (GFP-Cobl-like^1-457ΔKRAP1^) in the presence and absence of Ca^2+^, respectively (**D**). Arrowhead, increase of coimmunoprecipitated Flag-syndapin I with GFP-Cobl-like^1-457^ upon Ca^2+^. (**E**) Quantitation anti-syndapin I coimmunoprecipitation upon Ca^2+^ presence normalized to immunoprecipitated GFP-Cobl-like^1-457^ and GFP-Cobl-like^1-457ΔKRAP1^, respectively (as deviation from conditions without Ca^2+^). (**F,G**) Side-by-side comparison of syndapin I coimmunoprecipitations with GFP-Cobl-like^1-457^ and GFP-Cobl-like^1-457ΔKRAP1^ (**F**) and quantitative analysis thereof (**G**). Data, mean±absolute error. White line, lanes omitted from blot. (**H-J**) Functional analyses of the importance of Cobl-like’s CaM-regulated syndapin I binding site (KRAP1) by loss-of-function rescue experiments evaluating the indicated dendritic arbor parameters of developing neurons (transfection, DIV4; analysis, DIV5.5). Note that neither a Cobl-like mutant lacking the entire N terminal part (GFP-Cobl-like*^Δ1-412^) nor GFP-Cobl-like*^ΔKRAP1^ was able to rescue Cobl-like’s loss-of-function phenotypes. Data, **E**, bar/dot plot overlays with mean±SEM. Unpaired student’s t-test (**E**); mean±absolute error (**G**); mean±SEM; 1-way ANOVA+Tukey (**H-J**).

Strikingly, quantitative coimmunoprecipitation analyses unveiled a full abolishment of the about 50% increase of syndapin I interaction with the Cobl Homology domain of Cobl-like (Cobl-like^1-457^) upon Ca^2+^ addition when the first “KRAP” motif (KRAP1) was deleted (Cobl-like^1-457ΔKRAP1^; Cobl-like^1-457Δ59-69^) (**Figure 10C-E**). This insensitivity of Cobl-like^1-457ΔKRAP1^ to Ca^2+^/CaM signaling revealed that it was exclusively the first “KRAP” motif (aa59-69) that was regulated by Ca^2+^/CaM signaling.

Side-by-side analyses of GFP-Cobl-like^1-457^ and Cobl-like^1-457ΔKRAP1^ under Ca^2+^-free control conditions revealed that without Ca^2+^ Cobl-like^1-457^ and the corresponding ΔKRAP1 mutant thereof coimmunoprecipitated the same amount of syndapin I (**Figure 10F,G**). Thus, without Ca^2+^ and under the stringency of *in vivo* conditions, as reflected by coimmunoprecipitations, “KRAP” motif 1 seemed not to contribute to syndapin I complex formation but awaited activation by Ca^2+^/CaM signaling.

### The single Ca^2+^/CaM-regulated syndapin I binding site of Cobl-like is crucial for Cobl-like’s function in dendritic arbor formation

In line with the importance of the identified N terminal CaM binding site, also deletion of only the first, i.e. the Ca^2+^/CaM-regulated, syndapin I binding interface was as detrimental for Cobl-like’s critical functions in dendritic arborization as lacking the entire N terminal part of Cobl-like all together (GFP-Cobl-like*^Δ1-412^). Both GFP-Cobl-like*^Δ1-412^ and GFP-Cobl-like*^ΔKRAP1^ completely failed to rescue the Cobl-like loss-of-function phenotypes in dendritic arborization (**Figure 10H-J**).

Instead, cotransfections with Cobl-like RNAi in both cases merely led to dendritic morphologies identical to those of neurons deficient for Cobl-like. The dendritic branch points, terminal points and total dendritic length all remained significantly reduced in comparison to control neurons (scrambled RNAi/GFP) and did not differ from those of Cobl-like RNAi neurons (**Figure 10H-J**).

This complete failure to rescue any of the Cobl-like loss-of-function phenotypes in dendritic arborization demonstrated that the Ca^2+^/CaM-regulated “KRAP” motif 1 of Cobl-like is absolutely critical for Cobl-like’s functions in dendritic arbor formation.

Together, our analyses unveiled that the actin nucleator Cobl and its distant relative Cobl-like-each of them critical for dendritic arbor formation – in fact need to cooperate with each other in a syndapin-coordinated and Ca^2+^/CaM-regulated manner to bring about the complex morphology of hippocampal neurons (**Figure 11**).

**Figure 11.**
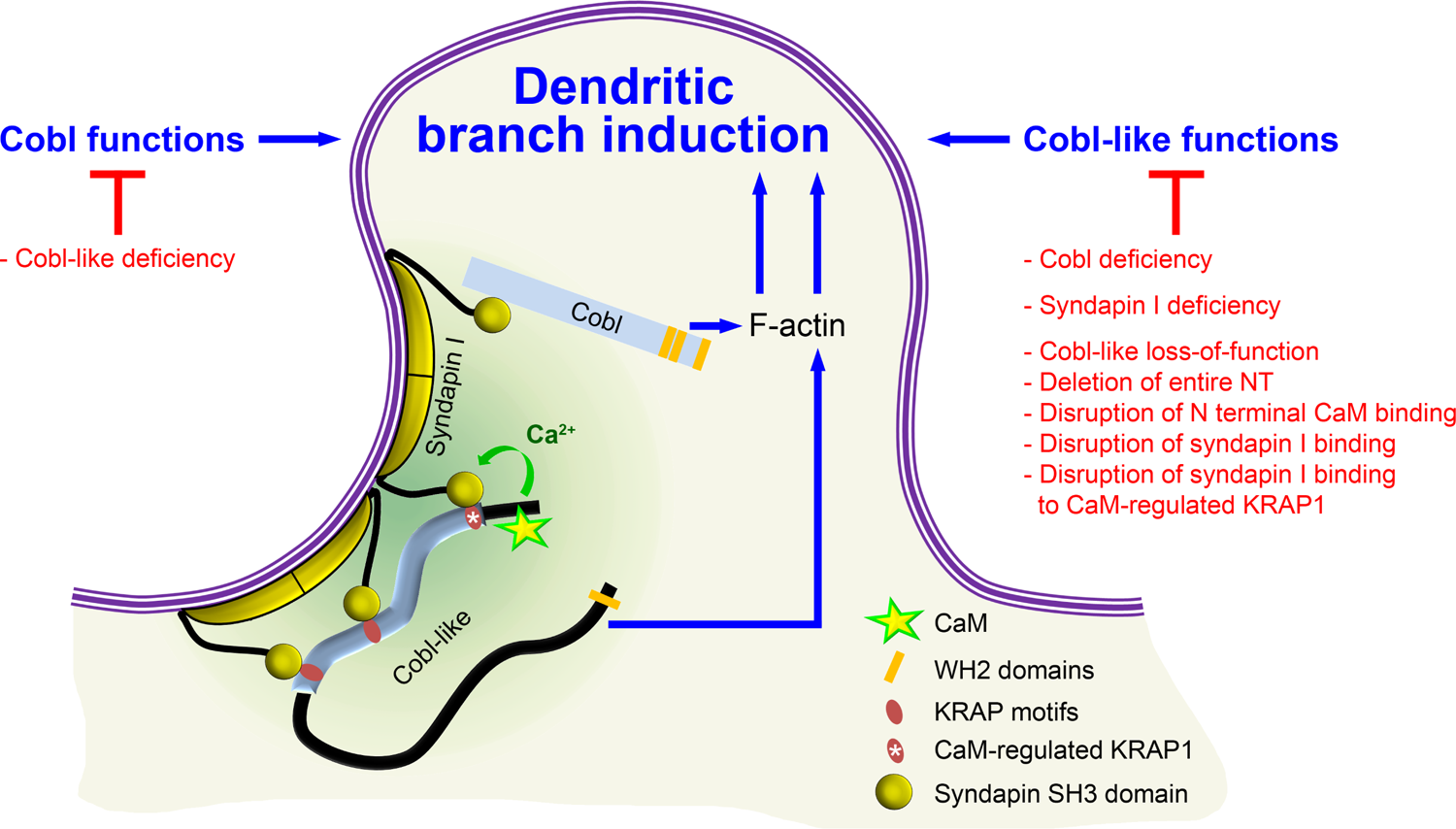
Model depicting how Cobl and Cobl-like functions in dendritic branch initiation are joined, coordinated and controlled. Cobl and Cobl-like functions are not only both critical for dendritic branch formation but both factors promoting the formation of actin filaments were found to act in an interdependent manner. The underlying mechanisms of coordination and control are depicted and include physical linkage of Cobl and Cobl-like by syndapin I forming dimers and multimeric clusters at the convexly bent membrane areas at the base of nascent branch sites. The newly identified interaction of syndapin I with Cobl-like is mediated by three independent KRAP motifs (red), the most N-terminal of which (marked by white asterisk) is regulated by a newly discovered CaM association to Cobl-like’s N terminus. All mechanistic aspects unveiled in this study are depicted in detail and the corresponding functional evaluations conducted are listed in brief. The WH2 domains of Cobl and Cobl-like are shown to indicate the C terminal domains of both proteins and their cytoskeletal functions.

## Discussion

Development of proper dendritic arbors of neuronal cells is key for the complex brains of vertebrates, as neuronal morphologies have direct consequences for brain organization patterns, cell-cell connectivity and information processing within neuronal networks. Here we show that this fundamental process is powered by the coordinated, strictly interdependent action of two components, which both promote the formation of actin filaments at the cell cortex, the actin nucleator Cobl (Ahuja et al., 2007) and its only distant relative Cobl-like (Izadi et al., 2018).

Cobl-like is already present in bilateria and considered as an evolutionary ancestor of the actin nucleator Cobl. Yet, we did not observe any redundant or additive functions in dendritic arborization of developing neurons. Instead, Cobl and Cobl-like both enriched at the same nascent dendritic branching sites and their functions were cooperative and each crucial for dendritic branch induction.

Our findings that Cobl-like interacts with the F-BAR domain protein syndapin I providing links to Cobl and that the syndapin I-binding N terminal part of Cobl-like is regulated by Ca^2+^/CaM signaling in a positive manner unveil the mechanisms of the striking functional interdependence of Cobl and Cobl-like (**Figure 11**).

The identified syndapin/Cobl-like interactions were mediated by three “KRAP” motif-containing regions in Cobl-like and the SH3 domain of syndapin I. Cobl-like’s “KRAP” motifs are highly conserved among each other and among different species (consensus, Kr+APxpP). They furthermore show similarity to those of Cobl (consensus, KrRAPpPP) (Schwintzer et al., 2011), as well as to other mapped syndapin I binding sites, such as RRQAPPPP in dynamin I (Anggono and Robinson, 2007), RKKAPPPPKR in ProSAP1/Shank2 (Schneider et al., 2014), and KKPPPAKPVIP in the glycine receptor beta subunit (del Pino et al., 2014).

Coprecipitation of endogenous syndapin I with Cobl-like from brain extracts, coimmunoprecipitations of endogenous Cobl-like and syndapin I from mouse brain lysates as well as visual proof of complex formation in intact cells underscore the *in vivo*-relevance of the Cobl-like/syndapin I interactions we identified.

Syndapin I/Cobl-like interactions clearly were of functional importance, as syndapin I deficiency completely suppressed Cobl-like-mediated dendritic arbor formation. In line, syndapin I accumulated together with Cobl-like at nascent dendritic branch sites and membrane-bound syndapin I clusters were found at convex membrane curvatures at the base of protrusions in developing neurons. With their topology changes in different directions, these membrane areas fit the structure of the membrane-binding F-BAR domain of syndapin I, which seems unique among the BAR protein superfamily (Qualmann et al., 2011) and shows overall curvature but also a strongly kinked tilde shape (Wang et al., 2009).

We furthermore demonstrated biochemically and in intact cells that Cobl-like and Cobl can physically be interconnected by syndapin I acting as a bridge. Physical interconnection of Cobl and Cobl-like by syndapin I provides a plausible molecular mechanism for the striking functional interdependence of Cobl and Cobl-like and is in line with syndapin I/Cobl interactions (Schwintzer et al., 2011) as well as with syndapin I’s F-BAR domain-mediated self-association ability (Kessels and Qualmann, 2006; Shimada et al., 2007; Wang et al., 2009) (**Figure 11**). This would leave the SH3 domain of each syndapin I free for recruiting effector proteins, for spatially organizing them at specific, curved membrane areas at nascent dendritic branch sites and for thereby coordinating their functions in dendritic branch induction.

Importantly, we found that the interlinkage of Cobl-like and Cobl was not static. The Cobl-like/syndapin I interaction was regulated by Ca^2+^ signals. This is well in line with the involvement of transient, local Ca^2+^ signals in dendritic arborization of developing neurons (Rajan & Cline 1998; Fink et al., 2003; Gaudilliere et al., 2004) and with membrane targeting and cytoskeletal functions of Cobl also being controlled by Ca^2+^/CaM (Hou et al., 2015). The additional Ca^2+^/CaM regulation of the Cobl-like/syndapin I interface would now provide another key regulatory mechanism right at the interlinking bridge between Cobl and Cobl-like.

The regulatory mechanism is based on an N terminal stretch of amino acids of Cobl-like proteins in front of the so-called Cobl Homology domain, which is absent in Cobl proteins. With “KRAP1”, Ca^2+^ signaling allowed for the modulation of specifically one out of three syndapin I binding sites of Cobl-like (**Figure 11**).

Cobl-like mutants, which either solely lacked the N terminal CaM binding site or the single Ca^2+^/CaM-regulated syndapin binding “KRAP1”, both failed to rescue Cobl-like loss-of-function phenotypes in dendritic branching. The N terminal CaM association regulating the “KRAP1”/syndapin I interaction and consistently also the “KRAP1” thus were critical for Cobl-like’s functions in dendritogenesis.

Our data clearly shows that dendritic arborization of developing neurons requires the Ca^2+^/CaM- and syndapin I-coordinated, joined action of Cobl and Cobl-like. In other cells and/or cellular processes, Cobl and Cobl-like also seem to have their independent, individual functions, such as the critical role of Cobl in F-actin formation right beneath the sensory apparatus of outer hair cells in the inner ear, the loss of which correlated with defects in pericentriolar material organization, in postnatal planar cell polarity refinement and in hearing (Haag et al., 2018). Further studies of *Cobl* KO mice unveiled an importance of Cobl for a specialized set of filaments interconnecting structural elements in the F-actin-rich terminal web of microvilli-decorated epithelial cells in the small intestine. However, there were no indications of an additional Cobl-like involvement (Beer et al., 2020). Also in proteomic analyses of myoblasts, which upon IGFN1 deficiency show altered G-to-F-actin ratios, only Cobl was identified but not Cobl-like (Cracknell et al., 2020).

Likewise, apart from dendritic branching of neurons studied here, there are no hints on any Cobl roles in functions that the *Cobl-like* gene has been linked to, such as diabetes and obesity (Mancina et al., 2013; Sharma et al., 2017). Cobl-like was also suggested as biomarker for different cancer types (Gordon et al., 2003; Gordon et al., 2009; Wang et al., 2013; Han et al., 2017; Plešingerová et al., 2017; Takayama et al., 2018), to be suppressed by Epstein-Barr virus infection (Gillman et al., 2018) and to be involved in B-cell development (Plešingerová et al., 2018) but there are no hints on Cobl roles in any of these processes.

The extension of very fine and elaborately branched cellular structures over hundreds of micrometers, as in dendritogenesis of neurons, certainly represents an extreme and rather special case of cellular morphogenesis. It is therefore well conceivable that a joined action of both Cobl and Cobl-like is required to promote actin filament formation at locally restricted sites to drive further branching. It currently seems plausible that Cobl and Cobl-like’s different actin filament formation mechanisms – spatial rearrangement of three actin monomers by the three WH2 domains of Cobl generating actin nuclei (Ahuja et al., 2007) *versus* use of the single WH2 domain of Cobl-like and the actin-binding cofactor Abp1 in a structurally not fully understood trans-mechanism (Izadi et al., 2018) – and their different modes of regulation by Ca^2+^ (Hou et al., 2015; Izadi et al., 2018) represent the distinct functions of Cobl and Cobl-like that have to be combined to power dendritic arborization. The recently identified Cobl regulation by PRMT2-mediated arginine methylation (Hou et al., 2018) may potentially also provide a unique aspect that needs to be integrated into the joined, interdependent function of Cobl and Cobl-like in dendritic branch induction.

Although to our knowledge, besides the here reported functions of Cobl, no information on functional cooperations of any actin nucleators is available for any actin cytoskeletal process in neuronal function or development, some initial studies on other actin nucleators also hint towards more interactive roles than initially thought. The actin nucleator Spire works with formin 2 in *Drosophila* oocytes (Quinlan et al., 2007; Pfender et al., 2011; Montaville et al., 2014). The formin mDia1 was reported to synergize with the APC protein (Okada et al., 2010; Breitsprecher et al., 2012). mDia1 was furthermore very recently found to indirectly interact with the Arp2/3 complex functionally cooperating in cortical F-actin stiffening of mitotic HeLa cells (Cao et al., 2020). Furthermore, the actin nucleator JMY was found to interact with the Arp2/3 complex in *in vitro*-reconstitutions (Zuchero et al., 2009; Firat-Karalar et al., 2011). However, as JMY RNAi did not cause any statistically significant decline in cell migration (Zuchero et al., 2009; Firat-Karalar et al., 2011) – a process firmly established to involve the Arp2/3 complex - the functional importance of a putative cooperation of JMY with the Arp2/3 complex in the formation of actin filaments remains unclear.

While these initial observations and the in part apparently conflicting data show that we are only at the very beginning of identifying and understanding any cooperative functions of actin nucleators, the here studied cell biological process of dendritic branching highlights that clearly there are actin filament-driven cellular processes, which require the coordinated action of not only one but at least two effectors promoting the formation of F-actin. Our mechanistic and functional studies clearly demonstrate that with Cobl and Cobl-like shaping neurons into their complex morphologies involves regulated and physically coordinated interactions of different actin filament formation-promoting factors at the base of nascent dendritic protrusion sites.

## Material and Methods

### DNA constructs

Plasmids encoding for GFP-Cobl-like and parts thereof were described previously (Izadi et al., 2018) and generated by PCR using the EST clone UniProtID Q3UMF0 as template, respectively. GFP-Cobl-like^111-262^ and Cobl-like^1-111^ were generated by subcloning with the help of internal restriction sites. Additional Cobl-like deletion mutants were generated by combining the following forward primers, aa1 fw: 5’-AATTAGATCTATGGACCGCAGCGTCCCCGATCC-3’; aa261 fw: 5’-AAAGATCTGATATCAGCAGAGAG-3’; aa537 fw: 5’-AAAGATCTAAGGATCCTGATTCAGC-3’; aa740 fw: 5’-GCCTCAAGAGAATTCAGG-3’; aa376 fw: 5’-TTGAATTCTTAAACCATGATCGCTTC-3’; aa182 fw: 5’-TTAGATCTCCTACACCTATAATC-3’ with the following reverse primers, aa457 rv: 5’-AACTCGAGCCCGGGACCAAGGGAGC-3’; aa538 rv: 5’-TTCTCGAGTTAATCAGGATCCTTCTC-3’; aa741 rv: 5’-TCCTGAATTCTCTTGAGG-3’; aa540 rv: 5’-TTCTCGAGTTAATCAGGATCCTTCTC-3’; aa411 rv: 5’-GCAAGCTTGGTTTTCGAAGGTGG-3’; aa272 rv: 5’-AAGAATTCTCAGTTGTGTGATATTTG-3’; aa380 rv: 5’-TTGAATTCGAAGCGATCATGGTG-3’.

Cobl-like mutants lacking the N terminal CaM binding site Cobl-like^ΔCaM NT^ (Δaa11-45) were generated by fusing a PCR product (primers, aa1-10+46-51 fw: 5’-AAAGATCTATGGACCGCAGCGTCCCGGATCCCGTACCCAAGAATCACAAATTCCTG-3’ and aa741 rv: 5’-TCCTGAATTCTCTTGAGG-3’) with Cobl-like^740-1273^ using the internal EcoRI site (corresponding to aa740/741) to obtain the respective mutated full length protein.

Cobl-like mutants lacking only the first “KRAP” motif (Δaa59-69; ΔKRAP1) were generated by fusing a DNA fragment obtained by PCR using an RNAi-resistant Cobl-like construct (Izadi et al., 2018) as template and primers aa1 fw (5’-AATTAGATCTATGGACCGCAGCGTCCCCGATCC-3’) and aa58 rv (5’-TTAAGCTTGCTCTGACAAATATG-3’) with a second PCR product (primer aa70 fw 5’-TTAAGCTTGCCGAGACGAAGGGC-3’ and aa741 rev: 5’-TCCTGAATTCTCTTGAGG-3’) using a HindIII site. The resulting Cobl-like^1-741ΔKRAP1^ was used to either generate Cobl-like^1-457ΔKRAP1^ using an internal SmaI site corresponding to aa456/457 or fused to Cobl-like^740-1273^ to generate the respective full length Cobl-like mutant Cobl-like^ΔKRAP1^.

Cobl-like mutants lacking all three “KRAP” motifs were generated by PCR using primers aa70 fw 5’-TTAAGCTTGCCGAGACGAAGGGC-3’ and aa333 5’-TTAAGCTTTGCATCCGAGGGC-3’ rv and fusing the resulting PCR product with a second PCR product obtained by using primers aa413 fw 5’-TTAAGCTTCTGGCTCAGACTGATG-3’ and aa457 rv as well as with a PCR product resulting from the above described aa1 fw and aa58 rv primers to give rise to aa59-69+334-412 deletion construct (Cobl-like^1-457ΔKRAP)^. Using an internal SmaI restriction site, Cobl-like^1-457ΔKRAP^ was then fused to the more C terminal parts of Cobl-like to give rise to either Cobl-like^1-741ΔKRAP^ or Cobl-like^ΔKRAP^ mutants.

A Cobl-like deletion mutant lacking the N terminal Cobl Homology domain Cobl-like^Δ1-412^ was generated by fusing a PCR product (primers, aa413 fw: 5’-TTAAGCTTCTGGCTCAGACTGATG-3’ and aa741 rv: 5’-TCCTGAATTCTCTTGAGG-3’; with Cobl-like^740-1273^ to obtain Cobl-like^Δ1-412^. Plasmids encoding for GST fusion proteins of Cobl-like^1-411^ were generated by subcloning into pGEX-4T-2 (GE Healthcare). A plasmid encoding for TrxHis-Cobl-like^1-411^ was generated by PCR and subcloning into pET-32 (Novagen) (primers, aa1 fw: 5’-AATTAGATCTATGGACCGCAGCGTCCCCGATCC-3’; aa411 rv: 5’-GGGTCGACGGTTTTCGAAGGTGG-3’) using EcoRI and SalI sites.

The RNAi construct directed against mouse and rat Cobl-like coexpressing GFP (Cobl-like RNAi#1) and scrambled RNAi control were described before (Izadi et al., 2018; Pinyol et al., 2007). Additionally, Cobl-like RNAi and scrambled RNAi were inserted into a pRNAT vector coexpressing farnesylated mCherry (mCherryF) (pRNAT-mCherryF; Schneider et al., 2014). Plasmids for rescue attempts were built by replacing the GFP reporter by either RNAi-insensitive, GFP-Cobl-like (Cobl-like RNAi/Cobl-like* and scrambled RNAi/Cobl-like*; Izadi et al., 2018), or by mutant GFP-Cobl-like sequences generated based on Cobl-like*.

Plasmids encoding for GST-tagged SdpI full-length and SH3 domain (aa376-441), respectively, as well as for a P434L-mutated SH3 (SdpI^SH3mut^) were described previously (Qualmann et al, 1999; Qualmann and Kelly, 2000). An alternative syndapin I SH3 domain (aa378-441) was as described (Braun et al., 2005).

Plasmids encoding for Xpress-tagged syndapin I full-length, Flag-syndapin I and mitochondrially targeted syndapin I (Mito-mCherry-SdpI), syndapin I ΔSH3 (Mito-mCherry-SdpI^ΔSH3^) and mCherry (Mito-mCherry) were described by Qualmann et al. (1999), Qualmann and Kelly (2000), Kessels and Qualmann (2002), Braun et al. (2005) and Dharmalingam et al. (2009), respectively. Syndapin I-mRubyRFP was generated by subcloning syndapin I into a derivative of pEGFP-N1 containing mRubyRFP instead of GFP (Izadi et al., 2018). mCherryF-coexpressing SdpI RNAi (bp297-317; for validations see Dharmalingam et al., 2009) vector and the corresponding control pRNAT vector were described previously (Schneider et al., 2014).

GST-SdpII^SH3^ (SdpII-l, aa383-488) and GST-SdpIII^SH3^ (aa366-425) were described before (Qualmann and Kelly, 2000; Seemann et al., 2017). Flag-syndapin II-s was as described (Dharmalingam et al., 2009). Flag-syndapin III was generated by subcloning from GST-syndapin III (Braun et al., 2005) and insertion into pCMV-tag2B. GFP-Cobl, Flag-mCherry-Cobl and GFP-Cobl^1-713^ were described previously (Hou et al., 2015). Mito-GFP-Cobl^1-713^ was generated by subcloning the respective Cobl-encoding sequence into the mitochondrial targeting vector.

RNAi constructs against rat and mouse Cobl coexpressing farnesylated mCherry were generated by subcloning into pRNAT-mCherryF (Schneider et al., 2014). The control expressing scrambled RNAi and mCherryF were as described (Schneider et al., 2014). Correct cloning by PCR was verified by sequencing in all cases.

### Antibodies, reagents and proteins

Rabbit anti-Cobl-like antibodies were raised against a combination of two GST-Cobl-like fusion proteins (GST-Cobl-like^537-741^ and GST-Cobl-like^740-1015^) as described previously (Izadi et al., 2018). The antibodies were affinity-purified according to procedures described previously (Qualmann et al., 1999; Kessels et al., 2000). Anti-syndapin I and anti-syndapin III antibodies were described previously (Qualmann et al., 1999; Koch et al., 2011). Anti-GST and anti-TrxHis antibodies from guinea pig and rabbit were described before, too (Qualmann and Kelly, 2000; Braun et al., 2005; Schwintzer et al., 2011).

Polyclonal rabbit anti-GFP (ab290) was from Abcam. Monoclonal mouse anti-GFP antibodies (JL-8) were from Clontech (632381). Monoclonal mouse anti-Flag (M2) and anti-MAP2 (HM-2) antibodies as well as polyclonal rabbit anti-Flag antibodies (F3165, F7425) were from Sigma. Anti-Xpress antibodies were from Invitrogen (46-0528).

Secondary antibodies used included, Alexa Fluor488- and 568-labeled goat anti-guinea pig antibodies (A-11073, A-11075), Alexa Fluor488- and 568-labeled donkey anti-mouse antibodies (R37114, A10037), Alexa Fluor647- and 680-labeled goat anti-mouse antibodies (A-21236, A32723), Alexa Fluor488-labeled donkey anti-rabbit (A-21206), Alexa Fluor568-647- and 680-labeled goat anti-rabbit antibodies (A-11036, A-21245, A-21109) (ThermoFisher Scientific) as well as DyLight800-conjugated goat anti-rabbit and anti-mouse antibodies (ThermoFisher Scientific, 35571, 35521). Donkey anti-guinea pig antibodies coupled to IRDye680 and IRDye800, respectively, were from LI-COR Bioscience (925-32411 and 926-32411). Goat anti-rabbit, anti-guinea pig and anti-mouse-peroxidase antibodies were from Dianova (Jackson ImmunoResearch) (711-035-152, 106-036-003, 115-036-003). 10 nm gold-conjugated goat anti-guinea pig antibodies (EM.GAR10) for electron microscopical examinations of freeze-fractured samples were from BBI Solutions. MitoTracker Deep Red 633 was from Molecular Probes.

Sepharose 4B-coupled CaM was from GE Healthcare. GST- and TrxHis-tagged fusion proteins were purified from *E. coli* lysates using glutathione-agarose or -sepharose (Sigma; GenScript) and Talon metal affinity resin (Clontech), respectively, as described previously (Schwintzer et al., 2011; Qualmann & Kelly, 2000). After purification, fusion proteins were dialyzed against PBS, characterized by SDS-PAGE and snap-frozen and stored at −80°C.

Tag-free syndapin I and III were generated by expressing both proteins in the pGEX-6P vector (GE Healthcare) and cutting of the GST tag from purified proteins using PreScission protease (GE Healthcare) in 150 mM NaCl, 2 mM DTT and 20 mM HEPES pH 7.4 buffer overnight at 4°C (during dialysis after elution). Cleaved off GST and uncleaved GST fusion proteins were removed with glutathione-sepharose.

### *In vitro*-reconstitutions of direct protein-protein interactions

Direct protein/protein interactions were demonstrated by coprecipitation assays with combinations of recombinant TrxHis- and GST-tagged fusion proteins purified from *E. coli* and/or immobilized CaM (GE Healthcare), respectively, in 10 mM HEPES pH 7.4, 300 mM NaCl, 0.1 mM MgCl_2_, 1% (v/v) Triton X-100 supplemented with EDTA-free protease inhibitor cocktail as well as in some cases with 500 µM Ca^2+^ added.

Eluted proteins were analyzed by SDS-PAGE, transferred to PVDF membranes by either semi-dry or tank blotting and then subjected to immunodetection with anti-TrxHis and anti-GST antibodies. Primary antibodies were detected with fluorescent secondary antibodies using a Licor Odyssey System.

### Culture and transfection, and immunostaining of cells

Culturing of HEK293 and COS-7 cells and their transfection using TurboFect (ThermoFisher Scientific) as well as their immunolabeling was essentially done as described (Kessels et al., 2001; Haag et al., 2012). In reconstitutions and visualizations of protein complex formations at the surfaces of mitochondria in intact cells, mitochondria of COS-7 cells were stained with 0.2 µM MitoTracker Deep Red 633 in medium at 37°C for 30 min and cells were subsequently fixed with 4% (w/v) paraformaldehyde (PFA) for 7 min.

### Preparation of HEK293 cell lysates

HEK293 cells were washed with PBS 24-48 h after transfection, harvested and subjected to sonification for 10 seconds and/or lyzed by incubation in lysis buffer (10 mM HEPES pH 7.4, 0.1 mM MgCl_2_, 1 mM EGTA, 1% (v/v) Triton X-100) containing 150 mM NaCl and EDTA-free protease inhibitor Complete (Roche) for 20 to 30 min at 4°C (Kessels & Qualmann, 2006). Cell lysates were obtained as supernatants from centrifugations at 20000xg (20 min at 4°C).

### Coprecipitation of proteins from HEK293 cell lysates

Coprecipitation experiments with extracts from HEK293 cells expressing different GFP-fusion proteins were essentially performed as described before (Qualmann et al., 1999; Schwintzer et al., 2011). In brief, HEK293 cell lysates were incubated with purified, recombinant GST-fusion proteins immobilized on glutathione-sepharose beads (GenScript) for 3 h at 4°C. The reactions were then washed several times with lysis buffer containing 150 mM NaCl and EDTA-free protease inhibitor Complete. Bound protein complexes were subsequently eluted with 20 mM reduced glutathione, 120 mM NaCl, 50 mM Tris/HCl pH 8.0 (30 min RT) or obtained by boiling the beads in 4xSDS sample buffer.

For coprecipitations with CaM, HEK293 cell lysates were prepared in an EGTA-free lysis buffer containing 150 mM NaCl, EDTA-free protease inhibitor cocktail and 200 µM calpain I inhibitor. Cell lysates were supplemented with either 1 mM EGTA or 500 µM Ca^2+^ to obtain conditions without and with Ca^2+^, respectively. After incubation with 25 µl CaM-sepharose 4B (GE Healthcare) for 3 h at 4°C and washing, bound proteins were isolated by boiling in 4xSDS sample buffer. Lysates, supernatants and eluates were analyzed by immunoblotting.

Triple coprecipitations, i.e. the examinations of GST-Cobl-like^1-411^/syndapin/GFP-Cobl^1-713^ complexes with either syndapin I or syndapin III as bridging component were essentially performed as described above (lysis buffer containing 150 mM NaCl) except that the extracts from HEK293 cells expressing GFP-Cobl^1-713^ were not only incubated with immobilized GST-Cobl-like^1-411^ but also with tag-free syndapin I or syndapin III for 3 h. Bound proteins were eluted with 20 mM reduced glutathione, 120 mM NaCl, 50 mM Tris/HCl pH 8.0. Eluates and supernatants were separated by SDS-PAGE and analyzed by anti-syndapin I/III, anti-GST and anti-GFP immunoblotting.

### Coprecipitation of endogenous syndapin I from mouse brain lysates

For coprecipitation of endogenous syndapin I, brain lysates were prepared from mice sacrificed by cervical dislocation. Extracts were prepared using an Ultra Turrax homogenizer (Ika Ultra Turrax T5Fu; 20000 rpm, 10 s) in lysis buffer containing EDTA-free protease inhibitor Complete and supplemented with 100 mM NaCl and 200 µM calpain inhibitor I. After clearing the lysates from cell debris by centrifugation at 1000xg for 20 min, the supernatants were used to precipitate endogenous syndapin I by TrxHis-Cobl-like^1-411^ fusion proteins immobilized on Talon metal affinity resin. Bound proteins were eluted by boiling in sample buffer, separated by SDS-PAGE and analyzed by anti-syndapin I immunoblotting.

### Heterologous and quantitative coimmunoprecipitation analyses

Heterologous coimmunoprecipitations addressing Cobl-like/syndapin I, syndapin II and syndapin III interactions were done with lysates of HEK293 cells transfected with GFP-Cobl-like fusion proteins and GFP, respectively, in combination with Flag-tagged syndapins. The cell lysates were incubated with anti-Flag antibodies or non-immune IgGs in lysis buffer containing 100 mM NaCl and EDTA-free protease inhibitor Complete for 3 h at 4°C. Antibody-associated protein complexes were isolated by 2 h incubation with protein A-agarose (Santa Cruz Biotechnology) at 4°C. The immunoprecipitates were washed with lysis buffer containing 100 mM NaCl, eluted from the matrix by boiling in a mix of 2 M (final) urea and SDS-sample buffer and analyzed by immunoblotting.

Comparisons of GFP-Cobl-like^1-457^ and Cobl-like^1-457ΔKRAP1^ for their ability to associate with Flag-syndapin I were also done by anti-GFP immunoprecipitations from lysates of transfected HEK293 cells generated according to the procedure described above.

For quantitative evaluations of the regulation of Cobl-like/syndapin I complexes, anti-GFP immunoprecipitations of GFP-Cobl-like fusion proteins were done in the presence (2 µM CaCl_2_ added) and in the absence of Ca^2+^ (1 mM EGTA added), respectively.

The amounts of coimmunoprecipitated Flag-syndapin I were quantified based on the detection of fluorescent antibody signals using a Licor Odyssey System providing a linear, quantitative read-out over several orders of magnitude. Anti-syndapin I coimmunoprecipitation signals were normalized to the amounts of anti-GFP signal representing the immunoprecipitated material. This ensured that similar amounts of GFP-Cobl-like proteins were examined for their extent of Flag-syndapin I coimmunoprecipitation. Both fluorescence signals were detected on the same blot using the two different fluorescence channels of the Licor Odyssey System. Data were expressed as percent difference from Ca^2+^-free conditions.

### Endogenous coimmunoprecipitations from mouse brain extracts

Mice were sacrificed and the brain was cut into small pieces and homogenized in 10 mM HEPES pH 7.5, 30 mM NaCl, 0.1 mM MgCl_2_ and 1 mM EGTA with protease inhibitors. Afterwards, Triton X-100 was added (0.2% v/v final) and the homogenates were extracted for 1 h at 4-6°C. The samples were then centrifuged at 100000xg for 30 min at 4°C and the resulting supernatants (mouse brain extracts) were incubated with affinity-purified rabbit anti-Cobl-like antibodies and non-immune rabbit IgGs, respectively, bound to protein A agarose (preincubation at 4°C and washing with above buffer and 0.2% (v/v) Triton X-100 (CoIP buffer)). After 4 h of incubation at 6°C, the proteins bound to the protein A agarose were washed with ice-cold CoIP buffer, eluted with SDS sample buffer (100°C, 5 min) and analyzed by immunoblotting using anti-Cobl-like and anti-syndapin I antibodies.

### Microscopy

Images were recorded as z-stacks using a Zeiss AxioObserver.Z1 microscope (Zeiss) equipped with an ApoTome, Plan-Apochromat 100x/1.4, 63x/1.4, 40x/1.3 and 20x/0.5 objectives and an AxioCam MRm CCD camera (Zeiss). Digital images were recorded by ZEN2012. Image processing was done by Adobe Photoshop.

### Spinning disk live microscopy of developing neurons

Primary rat hippocampal neurons were transiently transfected using Lipofectamine 2000 at DIV6. For imaging, the culture medium was replaced by 20 mM HEPES pH 7.4, 140 mM NaCl, 0.8 mM MgCl_2_, 1.8 mM CaCl_2_, 5 mM KCl, 5 mM D-glucose (live imaging buffer) adjusted to isoosmolarity using a freezing point osmometer (Osmomat 3000; Gonotec).

Live imaging was conducted at 37°C 16-24 h after transfection employing an open coverslip holder, which was placed into a temperature- and CO_2_-controlled incubator built around a spinning disk microscope based on a motorized Axio Observer (Zeiss). The microscope was equipped with a spinning disk unit CSU-X1A 5000, 488 nm/100 mW OPSL laser and 561 nm/40 mW diode lasers as well as with a QuantEM 512SC EMCCD camera (Zeiss).

Images were taken as stacks of 7-17 images at Z-intervals of 0.31 µm depending on cellular morphology using a C-Apochromat objective (63x/1.20W Korr M27; Zeiss). The time intervals were set to 10 s. Exposure times of 50-200 ms and 3-12% laser power were used. Image processing was done using ZEN2012 and Adobe Photoshop software.

### Culturing, transfection and immunostaining of primary rat hippocampal neurons

Primary rat hippocampal neuronal cultures were prepared, maintained and transfected as described previously (Qualmann et al., 2004; Pinyol et al., 2007; Schwintzer et al., 2011). In brief, neurons prepared from hippocampi of E18 rats were seeded at densities of about 60000/well (24-well plate) and 200000/well (12-well plate), respectively. Cells were cultured in Neurobasal^TM^ medium containing 2 mM L-glutamine, 1x B27 and 1 µM/ml penicillin/streptomycin. The neurons were maintained at 37°C with 90% humidity and 5% CO_2_.

Transfections were done in antibiotic-free medium using 2 µl Lipofectamine2000 and 1 µg DNA per well in 24 well plates. After 4 h, the transfection medium was replaced by conditioned medium and neurons were cultured further. All analyses were done with several independent neuronal preparations.

Fixation was done in 4% (w/v) PFA in PBS pH 7.4 at RT for 5 min. Permeabilization and blocking were done with 10% (v/v) horse serum, 5% (w/v) BSA in PBS with 0.2% (v/v) Triton X-100 (blocking solution). Antibody incubations were done in the same buffer without Triton X-100 according to Kessels et al. (2001) and Pinyol et al. (2007). In brief, neurons were incubated with primary antibodies for 1 h at RT and washed three times with blocking solution. Afterwards, they were incubated with secondary antibodies (1 h, RT). Finally, the coverslips were washed with blocking solution, PBS and water and mounted onto coverslips using Moviol.

### Quantitative analyses of dendrites of primary hippocampal neurons

For loss-of-function analyses and the corresponding rescue experiments, as well as for suppressions of Cobl-like overexpression phenotypes, DIV4 hippocampal neurons were transfected with RNAi and control vectors, respectively, and fixed and immunostained about 34 h later (DIV5.5). 2-6 independent coverslips per condition per assay and neurons of at least 2 independent neuronal preparations were analyzed based on the anti-MAP2 immunostaining of transfected neurons.

Transfected neurons were sampled systematically on each coverslip. Morphometric measurements were based on anti-MAP2 immunolabeling of transfected neurons. Using Imaris 7.6 software, the number of dendritic branching points, dendritic terminal points, and dendritic filament length were determined and Sholl analyses (Sholl, 1953) were conducted.

For each neuron, a “filament” (morphological trace) was drawn by Imaris 7.6 software using the following settings: Largest diameter, cell body diameter; thinnest diameter, 0.2 µm; start seed point, 1.5 x of cell body diameter; disconnected points, 2 µm; minimum segment size, 10 µm.

Immunopositive areas that were erroneously spliced by Imaris or protrusions belonging to different cells, as well as filament branch points that the software erroneously placed inside of the cell body were manually removed from the filament. Parameters determined were saved as Excel files and subjected to statistical significance calculations using GraphPad Prism5 and Prism6 software (RRID:SCR_002798).

### Freeze-fracturing and immunogold labeling

Hippocampal neurons were grown for 7 days on poly-D-lysine-coated sapphire disks (diameter 4 mm; Rudolf Brügger, Swiss Micro Technology) in 24 well plates, washed with PBS and subjected to ultrarapid freezing (4000 K/s) as well as to freeze-fracturing as described for mature neurons (Schneider et al, 2014).

Freeze-fracturing of developing neurons led to low yields of rather fragile replica of inner and outer membrane leaflets, which, however, were preserved during subsequent washing, blocking and incubation (Wolf et al., 2019). Replica were incubated with guinea pig anti-syndapin I antibodies (1:50; overnight, 4°C) and 10 nm gold-conjugated secondary antibodies as described for mature neurons (Schneider et al., 2014).

Controls addressing the specificity of anti-syndapin I labeling of freeze-fracture replica included evaluations of labeling at the E-face (almost no labeling) and quantitative analyses of syndapin I labeling densities at control surfaces not representing cellular membranes (low, unspecific immunogold labeling at a density of only 0.4/µm^2^). Further controls including secondary antibody controls and labeling of syndapin I KO material were described previously (Schneider et al., 2014).

Replica were collected and analyzed using transmission electron microscopy (TEM) and systematic grid explorations as described (Schneider et al, 2014; Seemann et al., 2017). Images were recorded digitally and processed by using Adobe Photoshop software. All analyses were done with two independent neuronal preparations.

Membrane areas with parallel membrane orientations (cylindrical) were distinguished from protrusive topologies, as established previously (Wolf et al., 2019). Anti-syndapin I immunogold labeling densities were determined using the complete area of the respective membrane topology on each image (measured by using ImageJ).

Anti-syndapin I cluster analyses were conducted using circular ROIs of 70 nm diameter. Density of clusters at cylindrical and protrusive membranes were calculated considering ≥3 anti-syndapin I labels per ROI as one cluster. Additionally, anti-syndapin I labeling being single, paired and clustered in 3 to 8 labels, respectively, was analyzed as percent of total anti-syndapin labeling.

### Statistical analyses and sample-size estimation

No explicit power analyses were used to compute and predefine required sample sizes. Instead, all neuronal analyses were conducted by systematic sampling of transfected cells across coverslips to avoid any bias. Morphometric analyses were then conducted by using IMARIS software.

All data were obtained from 2-5 independent neuronal preparations seeded onto several independent coverslips for each condition for transfection and immunostaining. For each condition, n numbers of individual neurons ranging from about 30 to 40 were aimed for to fully cover the biological variances of the cells. Higher n numbers yielded from the systematic sampling were accepted, too (e.g. see the control in **Figure 1E-G** (n=45) and in **Figure 2L-N** (65)). Lower n numbers were only accepted for the established Cobl overexpression phenotype (n=24) and the Cobl-like-mediated suppression if it (also n=24), as results were clear and sacrificing further rats for further primary neuron preparations could thus be avoided (**Figure 2L-N**).

Outliers or strongly scattering data reflect biological variance and were thus not excluded from the analyses. All n numbers are reported directly in the figures of the manuscript and all are numbers of independent biological samples (i.e. neurons) or biochemical assays, as additional replicates to minimize measurement errors were not required because the technical errors were small in relation to the biological/biochemical variances.

All quantitative biochemical data (**Figure 8E**, **Figure 10E** and **Figure 10G**) are provided as bar and dot plot overlays to report the individual raw data and the deviations of individual data points around the mean. Quantitative data represent mean±SEM throughout the manuscript. Exceptions are **Figure 10G**, in which data represents mean±absolute error (n=2; no difference), and **Figure 7C**, as the percent of total labeling shown in **Figure 7C** per definition has no error.

Normal data distribution and statistical significance were tested using GraphPad Prism 5 and Prism 6 software (SCR_002798). The statistical tests employed are reported in the respective figure legends. Dendritic arbor parameters (number of dendritic branch points, number of terminal points and total dendritic length) were analyzed for statistical significance employing 1-way ANOVA and Tukey post-test throughout.

All Sholl analyses were tested by 2*-*way ANOVA and Bonferroni post-test. Quantitative evaluation of syndapin I coimmunoprecipitations with Cobl-like^1-741^, Cobl-like^1-457^ and Cobl-like^1-457ΔKRAP1^ were analyzed by unpaired student’s t-test. Anti-syndapin I immunogold labeling densities at different surfaces of freeze-fractured replica of membranes of developing neurons were analyzed by 1-way ANOVA and the densities of anti-syndapin I clusters were analyzed by two-tailed Student’s t-test.

Statistical significances were marked by * *p* <0.05, ** *p* <0.01, *** *p* <0.001 and **** *p* <0.0001 throughout. In addition, the numbers of *p* values are reported directly in the figures. Note that for *p* <0.0001 (****) no values were provided by the software Prism 6, as the *p* values are too small.

## Acknowledgments

We thank A. Kreusch, B. Schade and K. Gluth for excellent technical support. This work was supported by *DFG* grants KE685/4-2 to MMK as well as QU116/6-2 and QU116/9-1 to BQ.

## Author contributions

M. Izadi, E. Seemann, Dirk Schlobinski and Lukas Schwintzer generated data. M. Izadi, E. Seemann, B. Qualmann and M. Kessels evaluated data. M. Izadi, B. Qualmann and M. Kessels designed the project and wrote the manuscript.

## Legends of Figure Supplements

**Figure 1–figure supplement 1.**
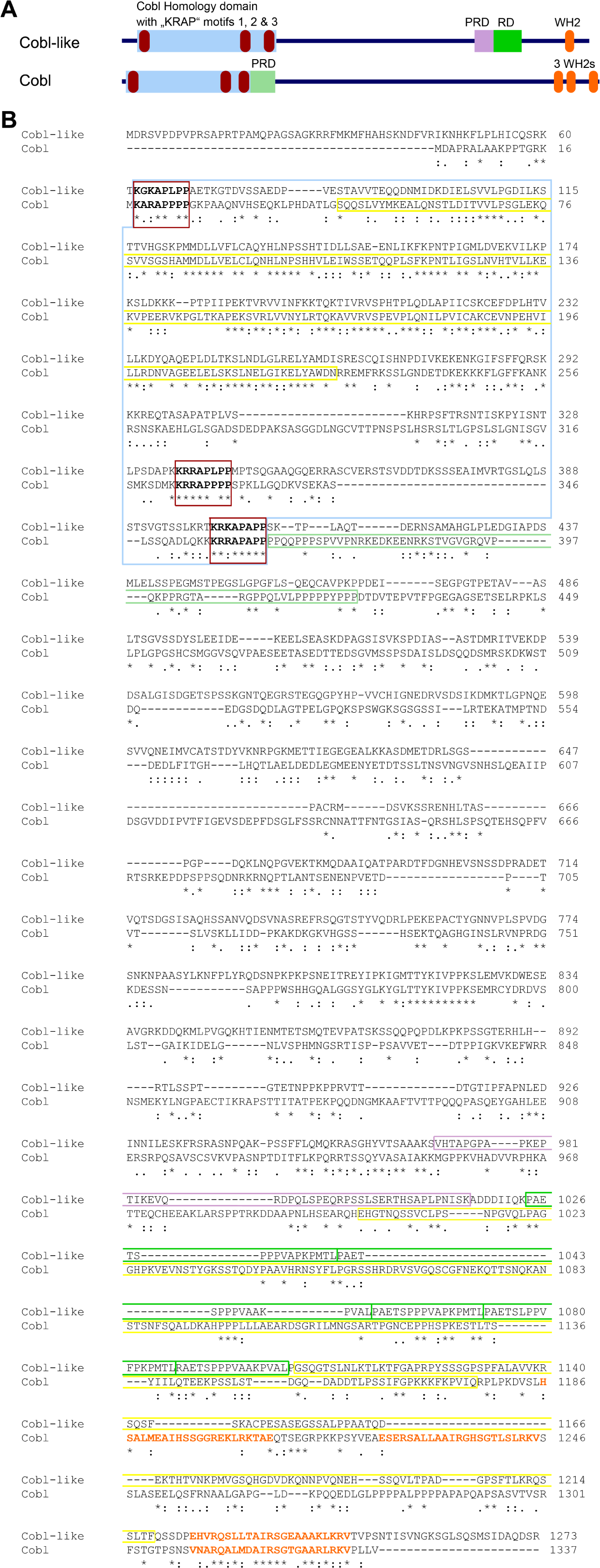
Comparison of domain structures and alignment of Cobl-like with the actin nucleator Cobl. (**A**) Scheme of Cobl-like with its domains in comparison to the actin nucleator Cobl with its domains. Abbreviations: PRD, proline-rich domain (PRD next to the Cobl Homology domain is an Abp1- and PRMT2-binding site and therefore colored in light green reminiscent of the RD, i.e. the Abp1-binding repeat domain in Cobl-like); WH2, WH2 domain. **(B)** Amino acid sequence alignment of murine Cobl-like (gi:74201419) and Cobl (gi:162135965) using Clustal Omega (https://www.ebi.ac.uk/Tools/msa/clustalo/). Marked are the Cobl Homology domains of both proteins (boxed in light blue) and the “KRAP” motifs (boxed in red) of both proteins. Furthermore, the single C terminal WH2 domain of Cobl-like and the three WH2 domains of Cobl are marked by orange letters. The CaM binding region of Cobl in the Cobl Homology domain and in the C terminal CaM binding regions in Cobl and Cobl-like are boxed in yellow, the Abp1- and PRMT2-binding PRD of Cobl is boxed in light green. The Abp1-binding repeat domain of Cobl-like is boxed in green and the proline-rich region located N terminal of the Abp1-binding area is boxed in purple. Note that although Cobl-like is considered as evolutionary ancestor of Cobl, the sequence conservation between both proteins in general is very low (*, identity; “:”, high similarity; “.”, moderate similarity). Even in the most-conserved part, the so-called Cobl Homology domain, the similarity is limited to the central core of this proposed domain and to the “KRAP” motifs.

**Figure 3–figure supplement 1.**
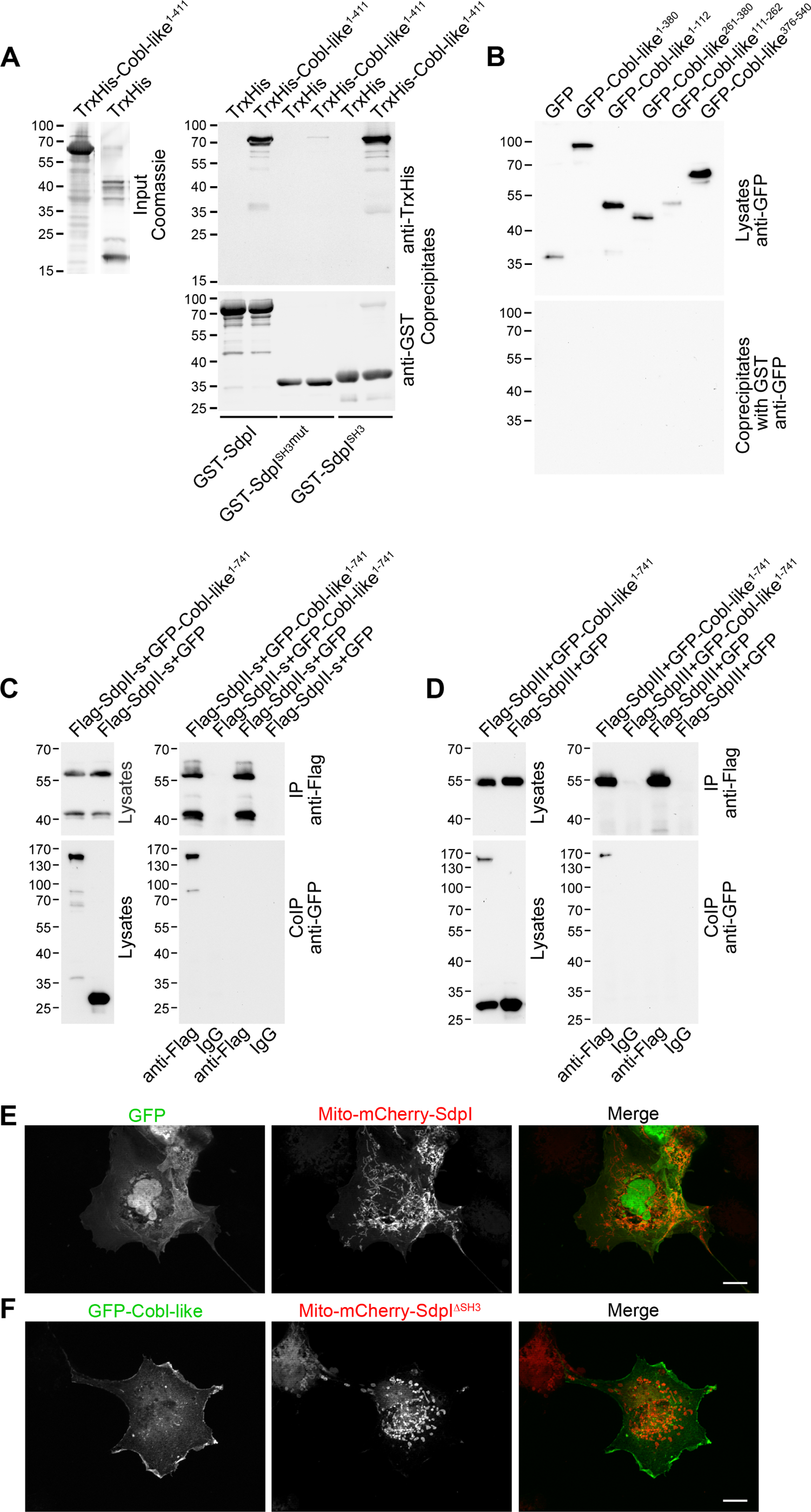
Cobl-like associates with syndapins. (**A**) Immunoblot analyses of a reconstitution of the association of TrxHis-Cobl-like^1-411^ with recombinant, purified GST-syndapin I and GST-syndapin I SH3 domain but not with a mutated syndapin I SH3 domain (P434L; SdpI^SH3mut^). (**B**) Anti-GFP immunoblotting of the GST controls of the syndapin I coprecipitation assays with Cobl-like deletion mutants shown in Figure 3E. (**C,D**) Immunoblotting analyses of specific coimmunoprecipitations of GFP-Cobl-like^1-741^ but not GFP with Flag-tagged syndapin II-s (short splice variant, SdpII-s) and syndapin III (SdpIII). (**E,F**) MIPs showing control experiments accompanying the protein complex reconstitutions between GFP-Cobl-like and GFP-Cobl-like^1-741^ and mitochondrially targeted syndapin I (Mito-mCherry-SdpI) shown in Figure 3G-I. (**E**) GFP is not recruited to Mito-mCherry-SdpI-enriched sites. (**F**) GFP-Cobl-like is not recruited to mitochondria when a Mito-mCherry-SdpI lacking the SH3 domain (Mito-mCherry-SdpI^ΔSH3^) is coexpressed. Bars, 10 µm.

**Figure 5–figure supplement 1.**
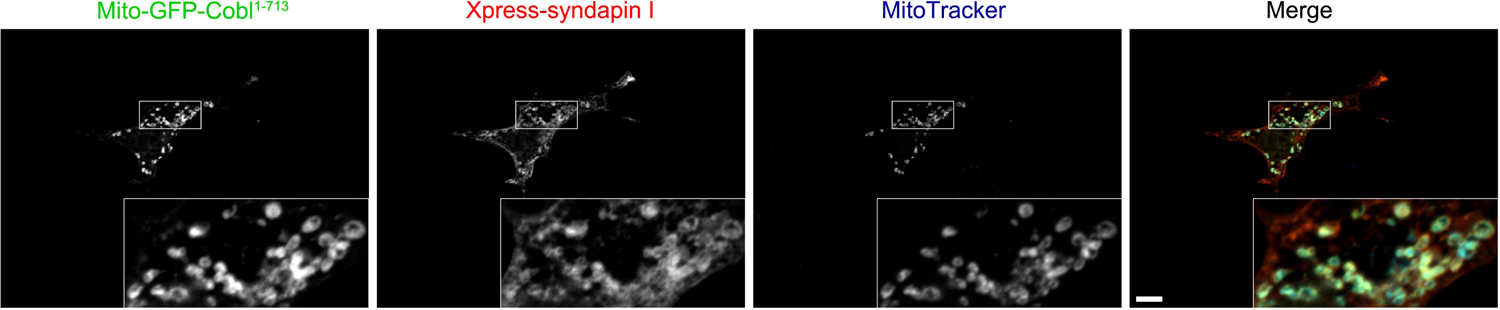
Recruitment of syndapin I to mitochondrial surfaces in intact cells by Mito-GFP-Cobl^1-713^. Reconstitution and visualization of Cobl/syndapin I protein complexes inside of intact COS-7 cells. Mitochondrially targeted Cobl^1-713^ (green in merge) recruits Xpress-tagged syndapin I (red in merge; anti-Xpress immunolabeling) to mitochondria (blue in merge; visualized by MitoTracker). Boxes mark the area presented as magnified inset (4fold enlargement). Colocalization of all three channels appears in whitish colors (white, beige, turquoise, rosé) in the merge. Bar, 10 µm.

**Figure 10–figure supplement 1.**
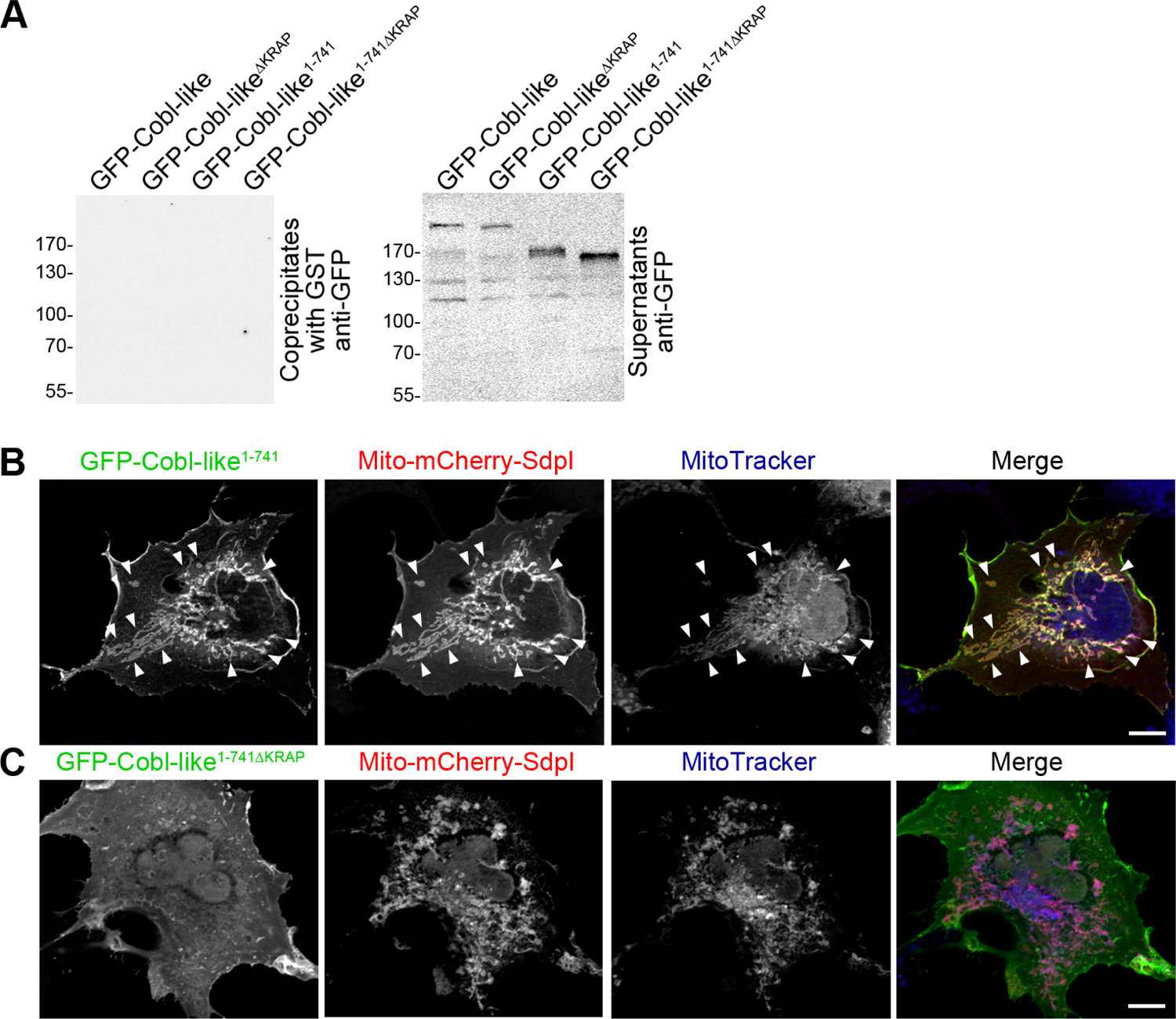
The three “KRAP” motifs are critical for Cobl-like’s association with syndapin I. (A) Anti-GFP immunoblotting of control experiments of the coprecipitation analyses shown in **Figure 10A** demonstrating that neither Cobl-like nor any of the Cobl-like mutants associated with GST. (**B,C**) MIPs of COS-7 cells transfected with Mito-mCherry-syndapin I and GFP-Cobl-like^1-741^ (B) and a GFP-Cobl-like mutant lacking the regions with the three “KRAP” motifs (GFP-Cobl-like^1-741ΔKRAP^) (**C**), respectively. Note the successful recruitment of GFP-Cobl-like^1-741^ to syndapin I-decorated mitochondria (colocalization of all three channels shown appears white in merge; examples are marked by arrowheads; **B**), whereas the corresponding GFP-Cobl-like^1-741ΔKRAP^ was not recruited to mitochondria (colocalization of only Mito-mCherry-syndapin I (red in merge) and MitoTracker (blue in merge) appears purple in merge; **C**). Bars, 10 µm.

